# Changes in Behavior and Salivary Serum Amyloid A in cows with Subclinical Mastitis

**DOI:** 10.1101/2021.01.10.426092

**Authors:** G. Caplen, S.D.E. Held

## Abstract

The aim of this study was to identify detailed changes in behavior, and in salivary serum amyloid A (**SAA**), associated with subclinical mastitis. This included standard sickness behaviors (which form part of an adaptive response to conserve energy, minimize heat loss and facilitate recovery following infection and inflammation) and lesser-studied behaviors, that are biologically predicted to change at low-level infection, and therefore particularly relevant for detecting subclinical mastitis (here labelled ‘luxury’ behaviors). SAA is a physiological marker of systemic inflammation, with levels in milk and serum already known to increase during subclinical mastitis. We investigated whether the same was true for SAA in cow saliva. Data were collected for n = 34 commercial barn-housed dairy cows, tested in 17 matched-pairs. Each pair comprised a cow with subclinical mastitis (**SCM)** and a healthy control (**CTRL**), identified using somatic cell count (**SCC**) (SCM: SCC >200 x1000 cells/ml; CTRL: SCC <100 x1000 cells/ml). SCM cows were selected for study ad-hoc, at which point they were paired with a CTRL cow, based upon parity and calving date; consequently, the full data set was accrued over several months. Data were collected for each pair over 3 days: SCC (l4:00-l5:00h) Day 1; behavior (24h from 00:00h) Day 2; salivary serum amyloid-A (SAA) Day 3. We report, for the first time, that an increase in salivary SAA occurs during subclinical mastitis; SAA was higher in SCM cows and demonstrated a positive (weak) correlation with SCC. The behavioral comparisons revealed that SCM cows had reductions in activity (behavioral transitions and distance moved), social exploration, social reactivity (here: likelihood to move away/be displaced following receipt of agonism), performance of social grooming and head butts, and the receipt of agonistic non-contact challenges. In addition, SCM cows received more head swipes, and spent a greater proportion of time lying with their head on their flank than CTRL cows. SCM cows also displayed an altered feeding pattern; they spent a greater proportion of feeding time in direct contact with two conspecifics, and a lower proportion of feeding time at self-locking feed barriers, than CTRL cows. Behavioral measures were found to correlate, albeit loosely, with serum SAA in a direction consistent with predictions for sickness behavior. These included positive correlations with both lying duration and the receipt of all agonistic behavior, and negative correlations with feeding, drinking, the performance of all social and all agonistic behavior, and social reactivity. We conclude that changes in salivary SAA, social behavior, and activity offer potential in the detection of subclinical mastitis and recommend further investigation to substantiate and refine our findings.

## 1. INTRODUCTION

Clinical stages of infectious disease are typically easily identified by obvious physical symptoms and behavioral changes (so-called ‘sickness behaviors’; Hart, 1988). Subclinical infection, that is infection below the level of clinical detection, by definition, is more difficult to spot. However, based on the interactions between the immune and central nervous system that cause sickness behavior predictions can also be made about behavioral changes during subclinical infection (Dantzer, 2004). These behavioral changes can be used as early warning signs of disease (Weary et al. 2009; von Keyserlingk et al. 2010), and/or to identify chronic subclinical infection levels. Mastitis remains a major concern in dairy cows with serious negative effects on welfare and productivity (Petersson-Wolfe et al. 2018). At subclinical levels, inflammation is present in response to the infection and milk production drops, but no abnormalities in the gland or milk are visible (Sordillo et al., 1997). It is therefore important for infection at any level to be identified and treated as soon as possible. Our main aim was to identify such changes in cows based on a detailed behavioral comparison between individuals with spontaneously occurring subclinical mastitis and healthy controls.

Behaviors in healthy animals can be divided, on the basis of immediate survival benefits, into ‘core maintenance’ and ‘luxury’ (after e.g. Dawkins, 1990). Core behaviors have immediate, short-term survival benefits; examples include resting, feeding, and drinking. They, therefore, start to decline only at later disease stages (e.g. Littin et al., 2008) and have low sensitivity during early and low-level stages (e.g. Sepúlveda-Varas et al., 2014). In our comparison we included standard ‘core’ sickness behaviors, such as changes to feeding, previously shown to be associated with experimentally-induced and spontaneously-occurring clinical mastitis (Siivonen et al., 2011; Fogsgaard et al. 2012; Sepúlveda-Varas et al., 2016). Because of our focus on subclinical mastitis, we also included so-called ‘luxury’ behaviors, such as physical and social exploration, grooming and various social interactions. These ‘luxury’ behaviors have delayed, longer-term, benefits and are not essential for immediate survival, meaning that they are biologically predicted to change at lower levels of disease than other sickness behaviors due to a greater sensitivity to disease challenge (Littin et al. 2008; Weary et al. 2009), especially when energy resources are diverted to fighting infection (see also ‘low-resilience behaviors’ in Littin et al., 2008).

In cows, effective health monitoring is hampered by logistical difficulties associated with direct animal observations within large open barns, reductions in human interaction linked with the substantial uptake in robotic milking units, and the tendency of cattle (for several biological reasons) to display only subtle indicators of pain/weakness (Gleerup et al., 2015). Advances in image analysis now allow automated recognition of individuals within a herd (Andrew et al., 2020), and accurate identification of health-related abnormal behaviors, e.g. foot disease (Gu et al., 2017). The identification of behaviors associated with subclinical disease (prior to the development of, or without, clinical symptoms) may therefore find application in future diagnostic software algorithms targeted at early disease monitoring in dairy cows (see e.g. Wagner et al. 2020).

A second aim was to quantify levels of salivary serum amyloid A (**SAA**), a major acute phase protein (**APP**) in cows (Murata et al., 2004), with and without spontaneously occurring subclinical mastitis. APPs are non-specific inflammatory markers that fluctuate in response to infection. Increased SAA levels have been detected during subclinical mastitis within both serum and milk (Kovac et al., 2011; Kovacevi-Filipovic et al., 2012). Other studies have confirmed SAA presence in bovine saliva (Lecchi et al., 2012; Rahman et al., 2013), suggesting salivary SAA has the potential for use in non-invasive detection of infectious diseases, such as mastitis.

Raised systemic inflammation levels, whether during acute infection or as chronic inflammatory states, are predicted to be accompanied by a feeling of sickness or ‘malaise’ in animals, as they are in humans (Dantzer et al., 2008; Weary et al. 2009). De Boyer Des Roches et al. (2017) reported correlations between (serum) SAA and several pain indicators before and after experimentally induced intra-mammary challenge with *E. coli*, including behavioral measures of attentiveness to surroundings. This suggested a direct link between serum SAA, infection levels and some sickness behavior in mastitic cows. A non-invasive means of monitoring systemic inflammation would also benefit future investigations into the impact of spontaneously occurring infections, such as mastitis, on cow welfare. In our study we tested for associations between salivary SAA levels and SCC (a standard measure of mastitis severity), and for associations between these physiological measures and behavior.

In summary, the purpose of this study was to identify differences in behavior and salivary SAA associated with subclinical mastitis. To this end we: (a) compared the behavior and salivary SAA of cows with subclinical mastitis with that of matched healthy individuals and, (b) correlated behavioral variables with both SCC and salivary SAA. We predicted that salivary SAA would be higher in cows with subclinical mastitis than in matched healthy controls. We also predicted that luxury behaviors, here including a range of social behaviors, would decrease with subclinical mastitis, and that differences detected would correlate negatively with the physiological measures (SCC and salivary SAA).

## 2. MATERIALS AND METHODS

### 2.1. Ethics Statement

The study was conducted between October 2017 and February 2018 at Bristol Veterinary School dairy farm. The experimental procedures were approved by the Animal Welfare and Ethical Review Board at the University of Bristol and conducted under University Investigation Number UB/17/061 ‘Behavioural markers of subclinical disease in dairy cows’.

### 2.2. Animals

Focal cows (n = 34) were part of an indoor commercial Holstein-Friesian dairy herd (n = 200) and resided within the low milk-yield group (approx. n = 80 cows) at the time of the study; having been part of the group for at least one month prior to data collection they were well-established within the social dominance hierarchy. Low yield animals were studied as they were deemed to be of low risk for having, or developing, subclinical metritis or ketosis during the trial. Cows were housed within a free-stall barn containing 93 lying cubicles (1.2 x 2.4m) with sand bedding, three stainless steel tip-over drinking troughs, a swinging brush (DeLaval), and automatic floor scrapers. Figure 1 shows the lay-out and relative positioning of resources within the pen. Cows were milked three times daily (at 06:00, 14:00 and 22:00h) and fed a total mixed ration once daily (06:00h).

**Figure 1:**
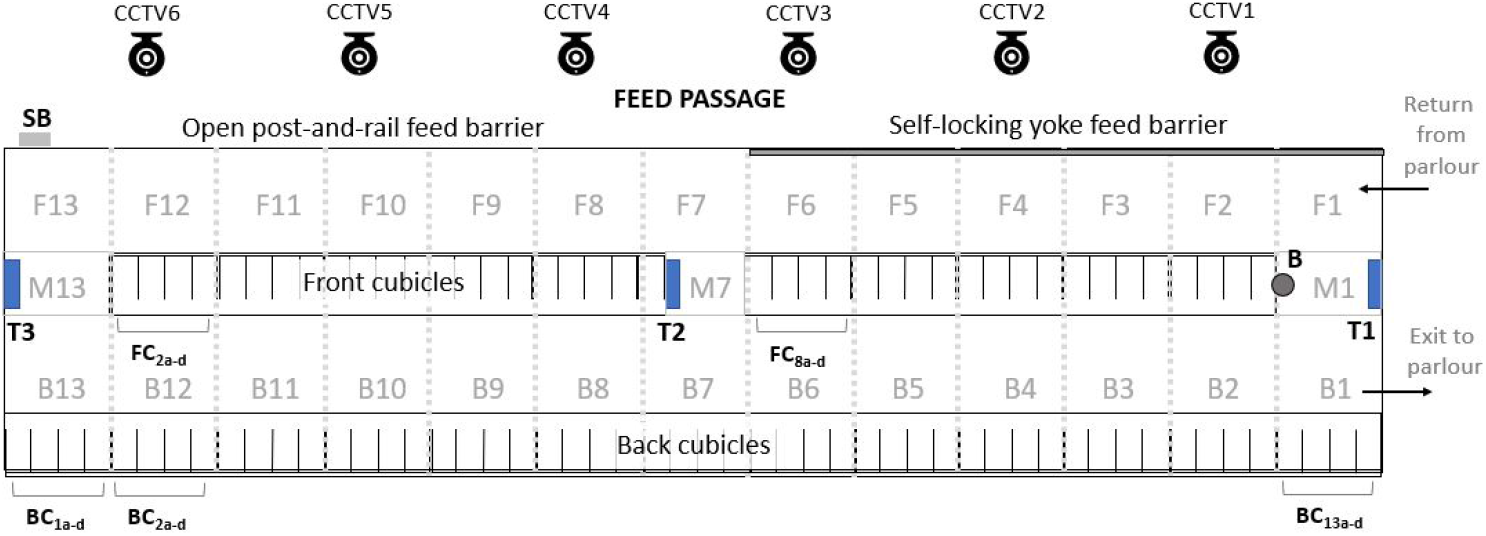
Plan of the home pen including CCTV camera position and virtual division of floor-space (F1-13, M1/7/13, B1-13) for logging cow position. SB = salt bin, T1-3 = water troughs, B = rotating brush

Only clinically healthy non-lame cows (mobility score ≤1; AHDB, 2015) without physical symptoms of mastitis were selected for inclusion in the study. Clinical health was assessed using individual health records and visual inspection of each cow by the researcher and herdsman. Data were collected concurrently from cows in matched pairs, each comprising a cow with subclinical mastitis (**SCM**) and a clinically healthy control (**CTRL**). Matching was performed on the basis of the following potential confounds: parity, pregnancy status (yes/no) and stage of pregnancy (days since insemination). Cows with a SCC of >200 (x1000 cells/ml) were classified as SCM (Madouasse et al., 2010), while cows with a SCC of <100 (x1000 cells/ml) were classified as CTRL. The 34 focal cows comprised 24 pregnant and 10 barren ones, 20 primiparous (n = 20) and 14 multiparous ones (of which two cows were at their 2^nd^ parity; two at 3^rd^; six at 4^th^; four at 5^th^). The time to expected calving date ranged from 55 - 236 days. While data from the two cows in a matched pair were collected at the same time, the data for 17 pairs took several months to collect due to low mastitis incidence within the herd at the time of the study.

### 2.3. Somatic Cell Count

Composite quarter milk samples were collected between 14:00-15:00h on Day 1. Although pairs of cows had data collected at different dates, for the purpose of our experimental design, all cows had SCC data collected on Day 1 and behavioral data on Day 2. Somatic cells were manually counted using a standard direct microscopic methodology (ISO 13366-1, 1997) following staining with Newman-Lampert stain solution: Levowitz-Weber modification (Newman’s Stain Solution: modified, 01375, Sigma-Aldrich).

### 2.4. Behavioral Measures

Each focal cow was fitted with a colored collar to facilitate individual recognition on Day 1. Two CCTV systems (N441L1T, Annke®, CA 91748, US), including six cameras recorded video footage from the entire low-yield pen. Continuous behavioral data were then coded retrospectively from video for each focal cow for 24h starting from 00:01h of Day 2 by a single experienced coder who was unaware of the health status of the cows at the time of scoring; these data made up the 24 h (‘**24h**’) data set. A second behavior data set was compiled for each cow using video recordings of the first 60 min following morning milking (‘**1hPostM1’**). This hour coincided with the peak feeding time of the day, since fresh feed was delivered to the feed corridor while the cows were in the parlour at morning milking. The start time of the 60 min observation period was specific to each focal cow, starting immediately following that cow’s re-entrance into the home pen from the parlour exit.

All behavioural measures are described in Table 1. Three broad categories of behavior were of interest: ‘core’ maintenance (including lying, feeding/drinking and activity), non-social ‘luxury’ behaviors (such as self-grooming and exploring the physical environment), and social ‘luxury’ behaviors. The last category was the most detailed, comprising non-agonistic (e.g. allogrooming, social exploration) and agonistic (e.g. head swipes, head butts) behaviors, in addition to social reactivity (here defined as the likelihood of moving away/being displaced following the receipt of an agonistic behavior).

**Table 1:**
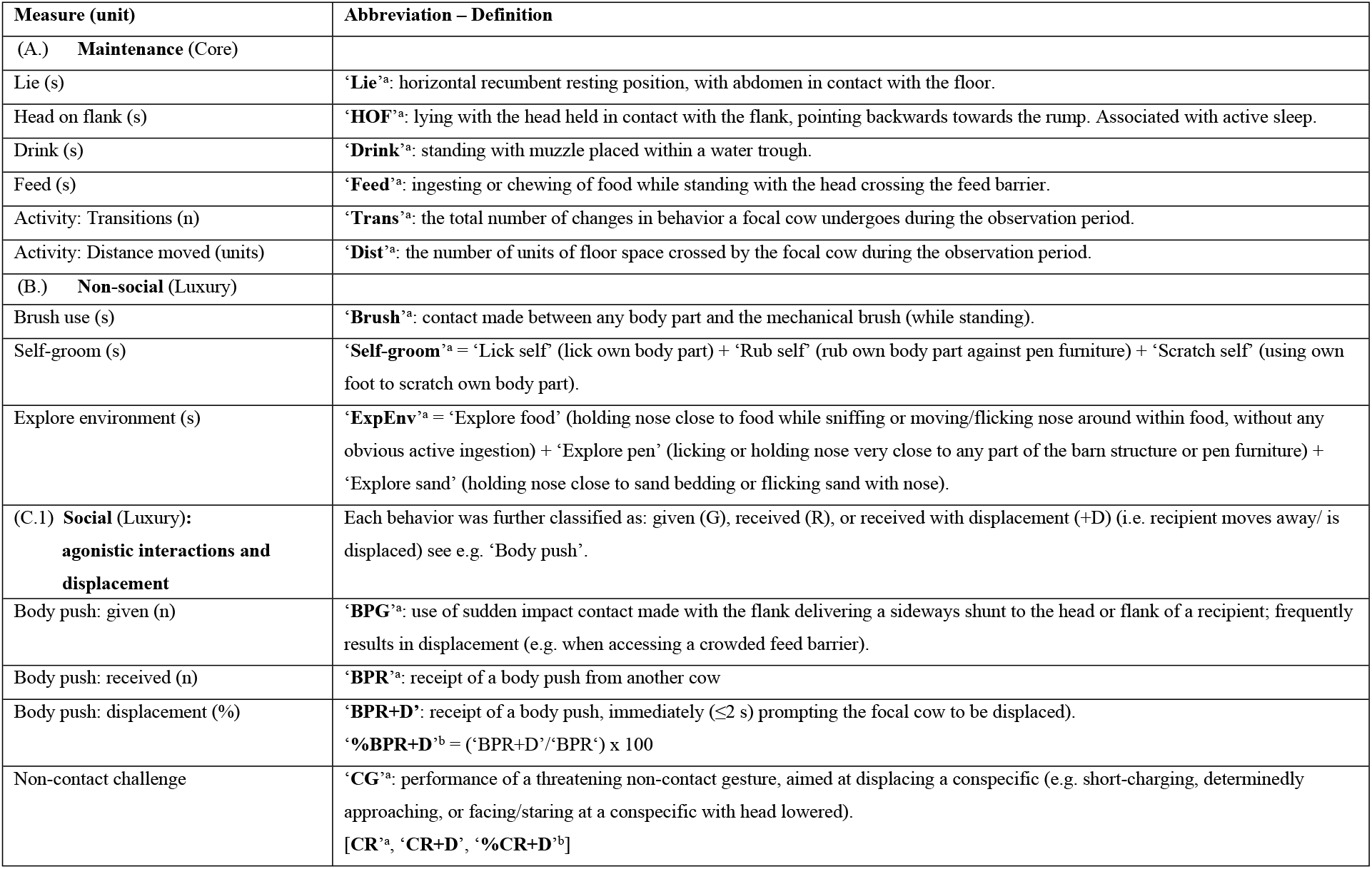
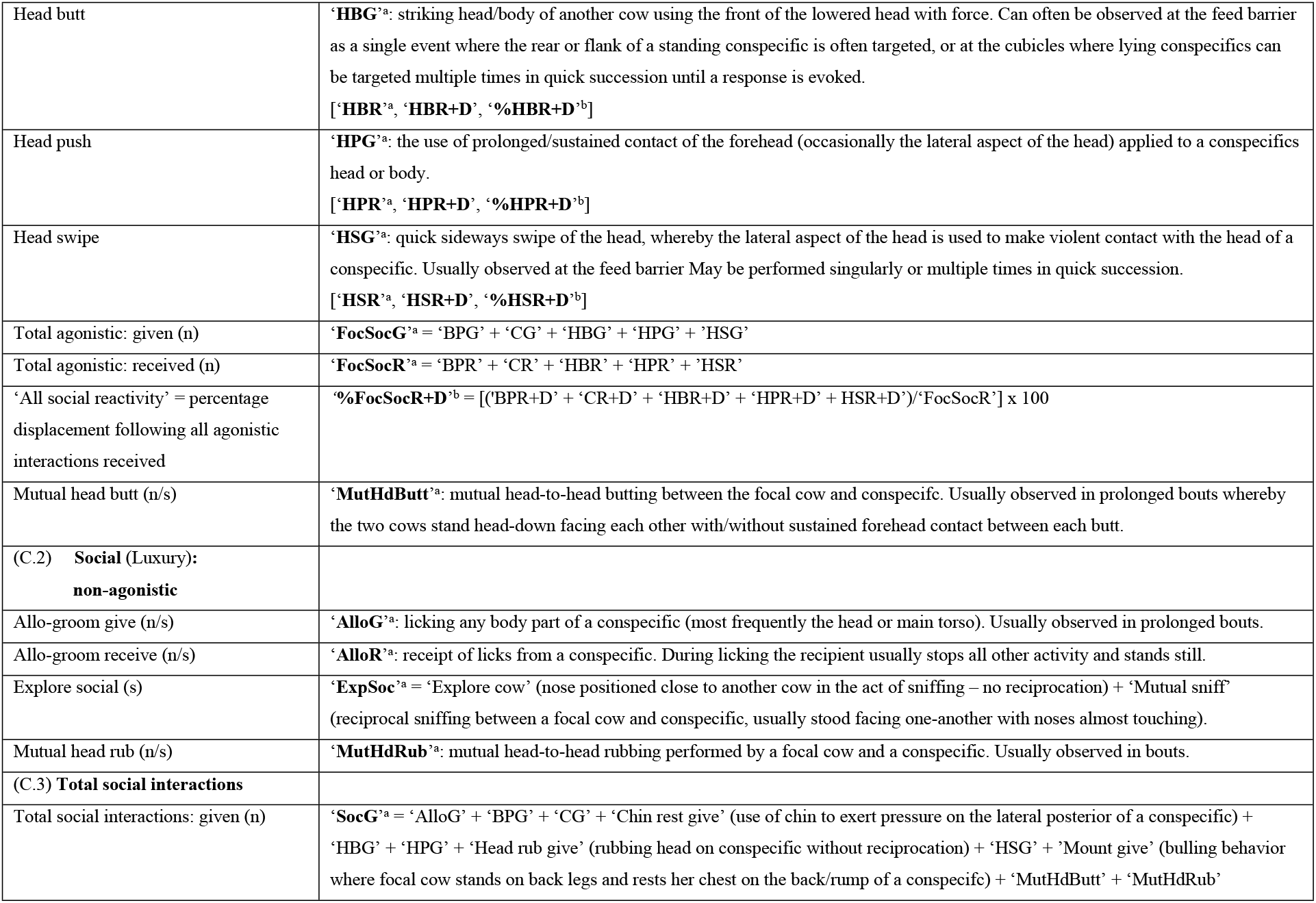
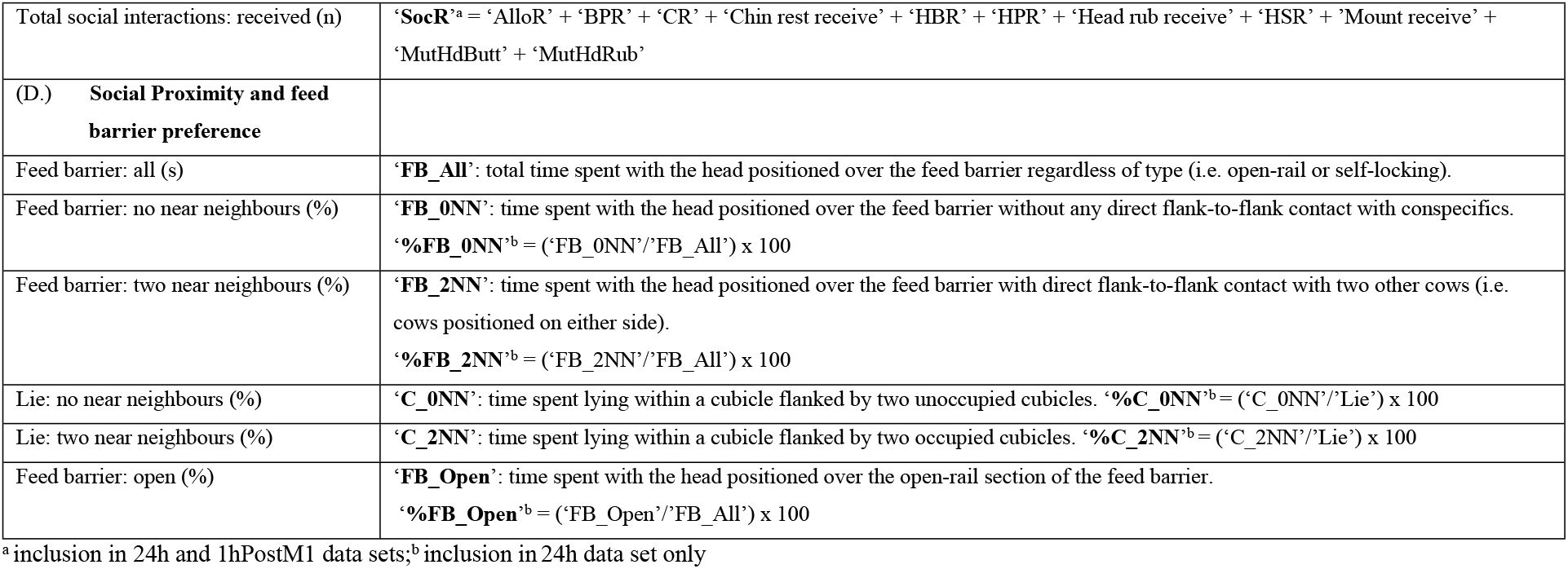
Cow behavioral measures used in the study

Total behavioral transitions (‘**Trans**’), a measure of activity, was calculated using all behaviors, including those coded, but not analysed individually (i.e. not listed in Table 1). These additional non-focal behaviors included eat sand, lick salt, paw sand, run, shake head, stand, and walk. Proximity was investigated using nearest neighbour (‘NN’) scores. When the focal cow was located at the feed barrier or resting in a cubicle, the number of other cows in immediate ‘proximity’ were scored as 0NN, 1NN or 2NN (Table 1). To enable an estimation of distance moved (‘**Dist**’), the pen floorspace was hypothetically subdivided into 29 units (each 4.8 m wide; see Figure 1) and the location of each cow was noted every five minutes throughout the 24h period. The number of floor units crossed during these intervals was then used to estimate distance moved.

Data were not available for 06:00, 14:00 or 22:00h as the cows were in the milking parlour or collecting yard during these periods. To account for differences in total time visible (i.e. due to variations in time spent within the parlour) data in the ‘24h’ data sets were standardised to either: number per hour visible (behavioral events) or seconds per hour visible (behavioral states); e.g. to standardise data from a cow that was visible for 20h, 42min and 12s within a 24h period data this data would be divided by 20.703.

### 2.5. Saliva Collection and SAA

Saliva was collected (Day 3) using a cotton swab (SalivaBio Children’s Swab, Item No. 5001.06, Salimetrics) and then immediately stored at −80°C prior to analysis. SAA was measured in saliva from 31 cows (saliva volumes from three cows were too small to be analysed), diluted 1:2, using a commercially available kit (Bovine Serum amyloid A protein ELISA Kit, EB0015, Finetest®, Wuhan Fine Biotech Co. Ltd.). To assess the suitability of the kit for use with saliva an assay validation was performed. To determine parallelism (linearity) a displacement curve, produced by double-diluting a pooled saliva sample with assay buffer, was compared to a standard curve. Percentage binding (as a percentage of that recorded for the zero standard) was calculated, in addition to the Log of the standard concentration (SAA standard) and the Log of the inverse of the dilution factor (saliva sample), e.g. 1:4 was transformed to Log(1/4). Parallelism was confirmed using a statistical test for the analysis of covariance (ANCOVA, SPSS). To measure assay accuracy the percentage recovery of exogenous SAA was calculated following the addition of 300 ng/ml SAA standard to a pooled saliva sample. Precision was assessed via intra- and inter-assay coefficients of variation (**CV**); the former was determined following the repeated measurement of aliquots of pooled saliva containing either high (quality control: **QC_high_**) or low (**QC_low_**) endogenous SAA within the same plate, while the later was determined following the assay of QC_high_ and QC_low_ samples in different plates.

### 2.6. Statistics

Following tests for normality (Shapiro-Wilk analysis), all behavioral measures, SAA and SCC were compared between CTRL and SCM using Paired samples t-test or Wilcoxon Signed-Rank tests (SPSS Statistics 24.0). Behavioral data from the continuous 24h data set and the data sub-set (60 min after the morning milking, ‘1hPostM1’) were separately analysed. Since the experimental design required the performance of multiple comparisons between measures there was an increased associated risk of Type I errors. Use of Bonferroni correction procedures has been highlighted as problematic (especially for animal behavioral studies, where sample sizes are often small) due to their tendency to increase Type II errors (Nakagawa, 2004). As an alternative to standard correction procedures we, therefore, calculated measures of observed (standardised) effect size in addition to p-values. Effect size measures the strength/magnitude of a relationship and, thereby, helps us to determine the strength of a statistical claim and whether a difference is real (i.e. it enables us to judge biological importance). Hedges’ g-value (Equations 1 and 2), also termed ‘Cohen’s d-value for paired samples’ (Hedges, 1981; Cohen, 1988; Nakagawa and Cuthill, 2007) and 95% confidence intervals (**CI**) for effect size (Equations 3 and 4), were calculated for all measures that met the assumptions of normality.

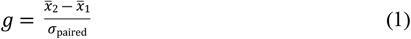

where

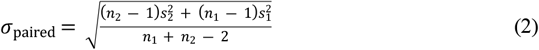

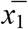 and 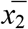 are the means of the two groups, *σ*_paired_ is the pooled standard deviation, *n* is the number of data points, and *s^2^* is the sample variance.

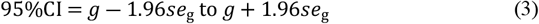

where

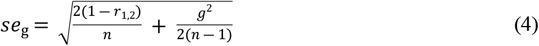

*se_g_* is the asymptotic standard error for the effect size, *n* = *n*_1_ = *n*_2_, and *r*_1,2_ is the correlation coefficient between the two groups. For all behavioral measures that did not meet the assumptions of normality, bootstrap effect size values (Hedges’ g-value with 95%CI, R = 2000) were computed using the software package ‘bootES’ (Gerlanc and Kirby, 2012; Kirby and Gerlanc, 2013) and R (Version 3.2.2., www.r-project.org/). Effect size statistics were interpreted as follows: (a) the size of the effect (based upon the estimated g-values: ≤0.39 = small, 0.40 - 0.79 = medium, ≥0.80 = large); (b) statistical significance (attributed to all measures where the associated 95%CI did not contain ‘0’) (Lee, 2016).

Interpretation of statistically non-significant p-values is possible using effect size confidence intervals in combination with the effect size (see Nakagawa & Foster, 2004). To identify those measures that were not statistically significantly different between SCM and CTL cows in the continuous 24h data set but could yet be biologically important, we used information from published studies to set broad, accepted, relative difference levels (**RDL%**, Table 2). Although these studies are not specific to subclinical mastitis, examples include the impact of clinical levels of (non-mastitic) infection and social status on behavior, it is assumed that they provide generous and relevant difference levels with which to compare our subclinical findings.

**Table 2:** Summary of the relative difference levels (RDL%) and their sources, used in this study to ascertain the existence of a biologically meaningful difference in behavioral measures between cows, with and without subclinical mastitis, based on effect size

| Behavioral Measure<sup>1</sup> | RDL% | Reference |
| --- | --- | --- |
| Agonistic social given: all categories | 10 | Sepulveda-Vares et al., 2016 |
| Agonistic social received: all categories | 10 | Neave et al., 2018 |
| %FB_0NN, %FB_2NN | 20 | Manson and Appleby, 1990 |
| %FB_Open | 10 | Huzzey et al., 2006 |
| AlloG | 30 | Galindo and Broom, 2002 |
| AlloR | 80 | Galindo and Broom, 2002; Hoonhout et al., 2017 |
| Brush | 30 | Mandel et al., 2017 |
| Self-groom | 20 | Fogsgaard et al., 2012 |
| Activity: Trans, Dist | 10 | King et al., 2018; Steensels et al., 2017 |
| Feed | 10 | Dollinger and Kaufmann, 2013 |
| Drink | 20 | Huzzey et al., 2007 |
| Lie | 10 | Toaff Rosenstein et al., 2016 |
| Other | 20 |  |

<sup>1</sup>%FB\_0NN = percentage of time at feed barrier with no near neighbors; %FB\_2NN = percentage of time at feed barrier with two near neighbors; %FB\_Open = percentage of time at open-rail feed barrier; AlloG = allogroom give; AlloR = allogroom receive; Brush = use of revolving brush; Self-groom = licking, rubbing, scratching self; Trans = behavioral transitions; Dist = distance moved; Lie = lying in cubicle.

For those measures where no relevant literature was available (‘Other’; Table 2), we used the average of the known RDLs (i.e. 20%; Table 2). For each measure, relative difference values (**RDV**) were then calculated using the RDL% and the respective mean value from the CTRL group. 95%CI_RDV_ were calculated using the confidence intervals from the effect size statistics and the (between-group) difference in means. In those cases where the 95%CI_RDV_ did not include the RDV, we conclude (with 95% confidence) that the current study showed no important biological effect for that measure; we refer to these as ‘biologically unimportant’. In cases where the 95%CI_RDV_ did include the RDV, we conclude that a difference was inconclusive but plausible; we refer to these as ‘biologically inconclusive’. For example, if CTRL cows performed more body pushes than the SCM cows, yet this difference failed to reach statistical significance (P≥0.10), using the p-value alone we would dismiss this behavior as being unaffected by subclinical inflammation. However, if the RDV for this behavior was within the 95%CI_RDV_ range (e.g. RDV = 0.08, 95%CI_RDV_ = −0.08 to 0.27) we would conclude that, although this effect is biologically inconclusive based upon our evidence, the difference may become significant given a larger sample size. Alternatively, if the RDV, in the above example, was 0.3, then we would conclude that the effect was biologically unimportant.

To test for correlations between physiological (SCC and SAA) and behavioral measures, we performed curve estimation regression statistics using the continuous 24h data set (SPSS: ANOVA, coefficient of determination) following tests for normality. Due to the small sample size, standard deviations for the behavioral measures were already large, so outliers (± 2SD) deemed to be atypically/excessively low or high were considered and removed prior to data analysis. Such outliers comprised a maximum of a single data point (i.e. one matched pair) for any measure.

## 3. RESULTS

### 3.1. Behavioral Differences over 24 hours (24h data set), and during 60 min immediately following first milking (1hPostM1 data subset)

#### 3.1.1 Luxury behavior: Social

##### (a) Total social interactions

In the 24h data, no significant differences were found in the total performance or receipt of social interactions(Table 3). The difference in the total performance of social interactions was classified as biologically unimportant in the context of this dataset (‘**SocG**’: RDV = 0.50, 95%CI_RDV_ = −0.26 to 0.43). Total receipt of social interactions, however, was biologically inconclusive on the basis of RDV confidence intervals (‘**SocR**’: RDV = 0.44, 95%CI_RDV_ = −0.21 to 0.64). This means that given a larger sample size, significant differences might become apparent in the latter measure; i.e. it may be possible to confirm that SCM cows do receive fewer social interactions. Indeed, during the 60 minutes following morning milking, SCM cows did receive significantly fewer social interactions (1hPostM1; Table 4).

**Table 3:**
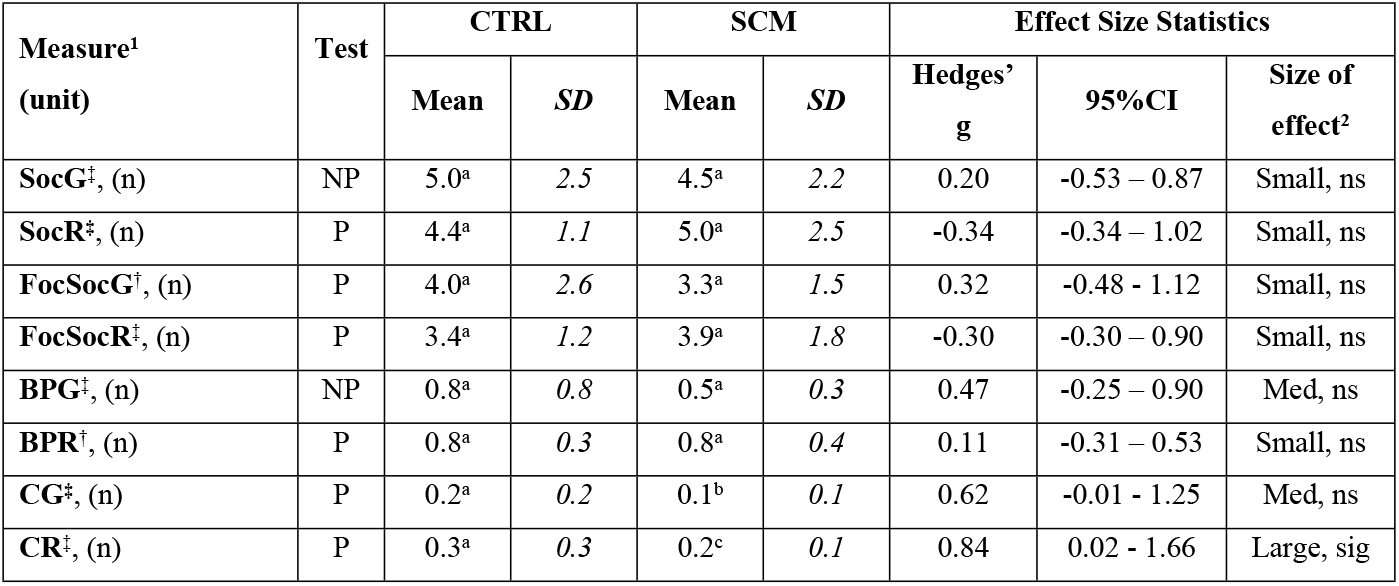
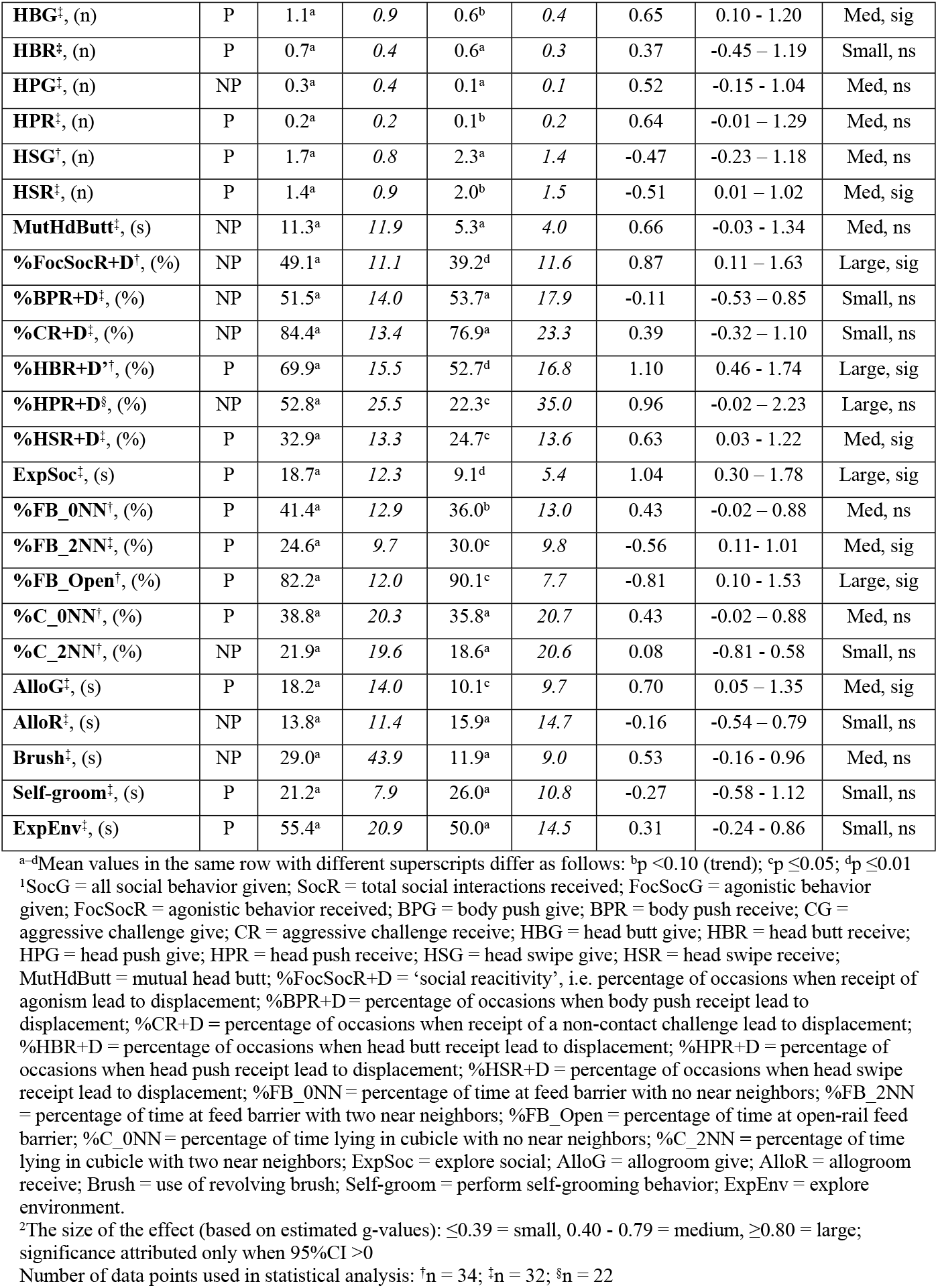
Measures of social and other ‘luxury’ behavior recorded from pair-matched cows with (SCM) or without (CTRL) sub-clinical mastitis over 24h: parametric data (P) analysed using paired t-test, non-parametric data (NP) analysed using Wilcoxon SR test. All units (except %) transformed to ‘per hour visible’.

| Measure<sup>1</sup><br>(unit) | Test | CTRL |  | SCM |  | Effect Size Statistics |  |  |
| --- | --- | --- | --- | --- | --- | --- | --- | --- |
|  |  | Mean | SD | Mean | SD | Hedges' g | 95%CI | Size of effect<sup>2</sup> |
| **SocG**<sup>‡</sup>, (n) | NP | 5.0<sup>a</sup> | 2.5 | 4.5<sup>a</sup> | 2.2 | 0.20 | -0.53 – 0.87 | Small, ns |
| **SocR**<sup>‡</sup>, (n) | P | 4.4<sup>a</sup> | 1.1 | 5.0<sup>a</sup> | 2.5 | -0.34 | -0.34 – 1.02 | Small, ns |
| **FocSocG**<sup>‡</sup>, (n) | P | 4.0<sup>a</sup> | 2.6 | 3.3<sup>a</sup> | 1.5 | 0.32 | -0.48 - 1.12 | Small, ns |
| **FocSocR**<sup>‡</sup>, (n) | P | 3.4<sup>a</sup> | 1.2 | 3.9<sup>a</sup> | 1.8 | -0.30 | -0.30 – 0.90 | Small, ns |
| **BPG**<sup>‡</sup>, (n) | NP | 0.8<sup>a</sup> | 0.8 | 0.5<sup>a</sup> | 0.3 | 0.47 | -0.25 – 0.90 | Med, ns |
| **BPR**<sup>‡</sup>, (n) | P | 0.8<sup>a</sup> | 0.3 | 0.8<sup>a</sup> | 0.4 | 0.11 | -0.31 – 0.53 | Small, ns |
| **CG**<sup>‡</sup>, (n) | P | 0.2<sup>a</sup> | 0.2 | 0.1<sup>b</sup> | 0.1 | 0.62 | -0.01 - 1.25 | Med, ns |
| **CR**<sup>‡</sup>, (n) | P | 0.3<sup>a</sup> | 0.3 | 0.2<sup>c</sup> | 0.1 | 0.84 | 0.02 - 1.66 | Large, sig |
| **HBG<sup>‡</sup>, (n)** | P | 1.1<sup>a</sup> | 0.9 | 0.6<sup>b</sup> | 0.4 | 0.65 | 0.10 - 1.20 | Med, sig |
| **HBR<sup>‡</sup>, (n)** | P | 0.7<sup>a</sup> | 0.4 | 0.6<sup>a</sup> | 0.3 | 0.37 | -0.45 – 1.19 | Small, ns |
| **HPG<sup>‡</sup>, (n)** | NP | 0.3<sup>a</sup> | 0.4 | 0.1<sup>a</sup> | 0.1 | 0.52 | -0.15 - 1.04 | Med, ns |
| **HPR<sup>‡</sup>, (n)** | P | 0.2<sup>a</sup> | 0.2 | 0.1<sup>b</sup> | 0.2 | 0.64 | -0.01 – 1.29 | Med, ns |
| **HSG<sup>‡</sup>, (n)** | P | 1.7<sup>a</sup> | 0.8 | 2.3<sup>a</sup> | 1.4 | -0.47 | -0.23 – 1.18 | Med, ns |
| **HSR<sup>‡</sup>, (n)** | P | 1.4<sup>a</sup> | 0.9 | 2.0<sup>b</sup> | 1.5 | -0.51 | 0.01 – 1.02 | Med, sig |
| **MutHdButt<sup>‡</sup>, (s)** | NP | 11.3<sup>a</sup> | 11.9 | 5.3<sup>a</sup> | 4.0 | 0.66 | -0.03 - 1.34 | Med, ns |
| **%FocSocR+D<sup>‡</sup>, (%)** | NP | 49.1<sup>a</sup> | 11.1 | 39.2<sup>d</sup> | 11.6 | 0.87 | 0.11 – 1.63 | Large, sig |
| **%BPR+D<sup>‡</sup>, (%)** | NP | 51.5<sup>a</sup> | 14.0 | 53.7<sup>a</sup> | 17.9 | -0.11 | -0.53 – 0.85 | Small, ns |
| **%CR+D<sup>‡</sup>, (%)** | NP | 84.4<sup>a</sup> | 13.4 | 76.9<sup>a</sup> | 23.3 | 0.39 | -0.32 – 1.10 | Small, ns |
| **%HBR+D<sup>‡</sup>, (%)** | P | 69.9<sup>a</sup> | 15.5 | 52.7<sup>d</sup> | 16.8 | 1.10 | 0.46 - 1.74 | Large, sig |
| **%HPR+D<sup>‡</sup>, (%)** | NP | 52.8<sup>a</sup> | 25.5 | 22.3<sup>c</sup> | 35.0 | 0.96 | -0.02 – 2.23 | Large, ns |
| **%HSR+D<sup>‡</sup>, (%)** | P | 32.9<sup>a</sup> | 13.3 | 24.7<sup>c</sup> | 13.6 | 0.63 | 0.03 - 1.22 | Med, sig |
| **ExpSoc<sup>‡</sup>, (s)** | P | 18.7<sup>a</sup> | 12.3 | 9.1<sup>d</sup> | 5.4 | 1.04 | 0.30 – 1.78 | Large, sig |
| **%FB_0NN<sup>‡</sup>, (%)** | P | 41.4<sup>a</sup> | 12.9 | 36.0<sup>b</sup> | 13.0 | 0.43 | -0.02 – 0.88 | Med, ns |
| **%FB_2NN<sup>‡</sup>, (%)** | P | 24.6<sup>a</sup> | 9.7 | 30.0<sup>c</sup> | 9.8 | -0.56 | 0.11 - 1.01 | Med, sig |
| **%FB_Open<sup>‡</sup>, (%)** | P | 82.2<sup>a</sup> | 12.0 | 90.1<sup>c</sup> | 7.7 | -0.81 | 0.10 - 1.53 | Large, sig |
| **%C_0NN<sup>‡</sup>, (%)** | P | 38.8<sup>a</sup> | 20.3 | 35.8<sup>a</sup> | 20.7 | 0.43 | -0.02 – 0.88 | Med, ns |
| **%C_2NN<sup>‡</sup>, (%)** | NP | 21.9<sup>a</sup> | 19.6 | 18.6<sup>a</sup> | 20.6 | 0.08 | -0.81 - 0.58 | Small, ns |
| **AlloG<sup>‡</sup>, (s)** | P | 18.2<sup>a</sup> | 14.0 | 10.1<sup>c</sup> | 9.7 | 0.70 | 0.05 – 1.35 | Med, sig |
| **AlloR<sup>‡</sup>, (s)** | NP | 13.8<sup>a</sup> | 11.4 | 15.9<sup>a</sup> | 14.7 | -0.16 | -0.54 – 0.79 | Small, ns |
| **Brush<sup>‡</sup>, (s)** | NP | 29.0<sup>a</sup> | 43.9 | 11.9<sup>a</sup> | 9.0 | 0.53 | -0.16 - 0.96 | Med, ns |
| **Self-groom<sup>‡</sup>, (s)** | P | 21.2<sup>a</sup> | 7.9 | 26.0<sup>a</sup> | 10.8 | -0.27 | -0.58 - 1.12 | Small, ns |
| **ExpEnv<sup>‡</sup>, (s)** | P | 55.4<sup>a</sup> | 20.9 | 50.0<sup>a</sup> | 14.5 | 0.31 | -0.24 - 0.86 | Small, ns |

**Table 4:** Differences in social and other ‘luxury’ behavior identified between pair-matched cows with (SCM) or without (CTRL) sub-clinical mastitis during 60 mins following morning milking (1hPostM1): parametric data (P) analysed using paired t-test, non-parametric data (NP) analysed using Wilcoxon SR test.

| Measure<sup>1</sup> | Test | CTRL |  | SCM |  | Effect Size Statistics |  |  |
| --- | --- | --- | --- | --- | --- | --- | --- | --- |
|  |  | Mean | SD | Mean | SD | Hedges' g | 95%CI | Size of effect<sup>2</sup> |
| **SocR**, (n) | NP | 12.1<sup>a</sup> | 7.43 | 6.6<sup>c</sup> | 3.92 | -0.91 | 0.17 – 1.51 | Large, sig |
| **FocSocG**, (n) | P | 10.7<sup>a</sup> | 9.0 | 5.8<sup>b</sup> | 4.7 | -0.70 | -0.05 – 1.45 | Med, ns |
| **FocSocR**, (n) | P | 4.7<sup>a</sup> | 2.6 | 2.5<sup>c</sup> | 1.9 | -0.99 | 0.04 – 1.93 | Large, sig |
| **BPR**, (n) | NP | 3.3<sup>a</sup> | 2.5 | 1.2<sup>c</sup> | 1.5 | -0.98 | 0.29 – 1.67 | Large, sig |
| **HBG**, (n) | NP | 1.7<sup>a</sup> | 2.4 | 0.5<sup>b</sup> | 0.6 | -0.65 | 0.10 – 1.09 | Med, sig |
| **HBR**, (n) | NP | 1.1<sup>a</sup> | 1.4 | 0.3<sup>b</sup> | 0.6 | -0.73 | 0.10 – 1.40 | Med, sig |
| **HPG**, (n) | NP | 0.9<sup>a</sup> | 1.3 | 0.1<sup>c</sup> | 0.3 | -0.84 | 0.24 – 1.38 | Large, sig |
| **ExpSoc**, (s) | NP | 21.2<sup>a</sup> | 28.8 | 6.5<sup>c</sup> | 7.6 | -0.68 | 0.06 – 1.07 | Med, sig |
| **AlloR**, (s) | NP | 37.6<sup>a</sup> | 66.1 | 1.9<sup>c</sup> | 3.2 | -0.74 | 0.44 – 1.11 | Med, sig |
| **Self-groom** (s) | P | 29.2<sup>a</sup> | 25.6 | 47.4<sup>b</sup> | 44.2 | 0.52 | 0.01 - 1.03 | Medium, sig |
| **ExpEnv**, (s) | P | 71.8<sup>a</sup> | 44.2 | 39.3<sup>c</sup> | 27.8 | -0.91 | 0.11 - 1.71 | Large, sig |

<sup>a-c</sup>Mean values in the same row with different superscripts differ as follows: <sup>b</sup>p < 0.10 (trend); <sup>c</sup>p ≤ 0.05.
<sup>1</sup>SocR = all social behavior received; FocSocG = agonistic behavior given; FocSocR = agonistic behavior received; BPR = body push receive; HBG = head butt give; HBR = head butt receive; HPG = head push give; ExpSoc = explore social; AlloR = allogroom receive; Self-groom = perform self-grooming behavior; ExpEnv = explore environment.
<sup>2</sup>The size of the effect (based upon estimated g-values): ≤ 0.39 = small, 0.40 - 0.79 = medium, ≥ 0.80 = large; significance attributed only when 95%CI > 0 (i.e. do not contain '0')

##### (b) Agonistic interactions

Broken down into separate interaction types, SCM cows performed significantly fewer head butts (‘**HBG**’: 24h; 1hPostM1) and head pushes (‘**HPG**’: 1hPostM1) than healthy controls (Tables 3 and 4). Non-significant measures of agonistic interactions given over 24h were classified as biologically inconclusive on the basis of RDV confidence intervals, again suggesting a larger sample might confirm an effect. These included body pushes (‘**BPG**’: RDV = 0.08, 95%CI_RDV_ = −0.08 to 0.27); non-contact challenges (‘**CG**’: RDV = 0.02, 95%CI_RDV_ = 0.00 to 0.15), head pushes (‘**HPG**’: RDV = 0.03, 95%CI_RDV_ = −0.03 to 0.18), and head swipes (‘**HSG**’: RDV = 0.17, 95%CI_RDV_ = −0.12 to 0.63) as well as total agonistic interactions (‘**FocSocG**’: RDV = 0.40, 95%CI_RDV_ = −0.31 to 0.73).

Over the 24h period, SCM cows received significantly fewer non-contact challenges than healthy cows, but more head swipes (Table 3). With the exception of body push received, which was classified as biologically unimportant (‘**BPR**’: RDV = 0.08, 95%CI_RDV_ = −0.01 - 0.02), all non-significant measures of agonistic interactions received over 24h were classified as biologically inconclusive on the basis of RDV confidence intervals, indicating that a larger sample might confirm an effect. This included head butts (‘**HBR**’: RDV = 0.07, 95%CI_RDV_ = −0.05 to 0.13), head pushes (‘**HPR**’: RDV = 0.02, 95%CI_RDV_ = −0.00 to 0.14) and mutual butting (‘**MutHdButt**’: RDV = 1.13 s, 95%CI_RDV_ = −0.19 to 8.03 s), as well as total agonistic interactions (‘**FocSocR**’: RDV = 0.34, 95%CI_RDV_ = −0.13 to 0.40).

During the 60 minutes following morning milking cows with subclinical mastitis received significantly fewer total agonistic interactions than healthy controls, and specifically fewer body pushes and head butts (1hPostM1; Table 4).

##### (c) Social reactivity, social exploration, social proximity, and feed barrier preference

Over the 24h period, cows with subclinical mastitis were also significantly less ‘socially reactive’ than healthy cows (Table 3), i.e. were less likely to move away/be displaced following the receipt of agonistic interactions (see Table 1). They performed significantly less social exploration than healthy controls (‘**ExpSoc**’) during both the 24h period, and the 60 minutes following morning milking (Tables 3 and 4), and spent a significantly greater proportion of their time at the open section of the feed barrier over the 24h period (Table 3). Cows with subclinical mastitis spent a significantly greater proportion of their time at the feed barrier flanked by two neighbours over the 24h period than healthy controls (Table 3). On the basis of RDV confidence intervals, non-significant measures of social proximity were classified as biologically unimportant within the context of this study. This included proportion of time at the feed barrier without neighbors (‘**%FB_0NN**’: RDV = 8.28%, 95%CI_RDV_ = −0.11 to 4.75%) and the proportion of time lying within cubicles with zero or two neighbors (‘**%C_0NN’**: RDV = 7.75%, 95%CI_RDV_ = −0.06 to 2.58%; ‘**%C_2NN**’: RDV: 4.38%, 95%CI_RDV_ = −2.69 to 1.93%).

##### (d) Allogrooming

Over the 24h period, SCM cows performed allogrooming significantly less than healthy controls (Table 3). Although receipt of allogrooming was not significantly different during 24h, and this measure was classified as biologically unimportant on the basis of RDV confidence intervals (‘**AlloR**’: RDV = 11.02 s, 95%CI_RDV_ = −1.14 to 1.67 s), during 1hPostM1 the SCM cows were allogroomed less than the healthy controls (Table 4).

#### 3.1.2 Luxury behavior: Non-social

##### (a) Self-grooming and brush use

In the 60 minutes following morning milking there was a tendency for SCM cows to perform more self-grooming than healthy controls (Table 4). For the 24h period, differences in self-grooming and brush use were not significant but classified as biologically inconclusive on the basis of confidence intervals for effect size differences (‘Brush’: RDV = 8.70 s, 95%CI_RDV_ = −2.74 to 16.45 s; ‘Self-Groom’: RDV = 4.23 s, 95%CI_RDV_ = −2.81 to 5.42 s).

##### (b) Environmental exploration

In the 60 minutes following morning milking SCM cows explored the environment significantly less than healthy controls (Table 4). This difference was not evident over the 24h period (Table 3), instead being classified as biologically unimportant in the context of this study, as based upon RDV confidence intervals (‘**ExpEnv**’: RDV = 11.08 s, 95%CI_RDV_ = −1.30 to 4.66 s).

#### 3.1.3 Core maintenance behavior

##### (a) Feeding, drinking, and lying

For both the 24h observations and the 60 minutes following morning milking, no significant differences emerged in time spent feeding (‘**Feed**’), drinking (‘**Drink**’), or lying (‘**Lie**’) (Tables 5 and 6), nor was subclinical mastitis considered to have biologically important (or even inconclusive) effects on these behaviors over 24h (‘Feed’: RDV = 82.39 s, 95%CI_RDV_ = −0.03 to 0.03 s; ‘Drink’: RDV = 6.86 s, 95%CI_RDV_ = −1.33 to 3.99 s; ‘Lie’: RDV = 200.19 s, 95%CI_RDV_ = −23.54 to 79.67 s). However, over the 24h period SCM cows spent significantly more time lying with their head on their flank than healthy controls (‘**HOF’,** Table 5).

**Table 5:**
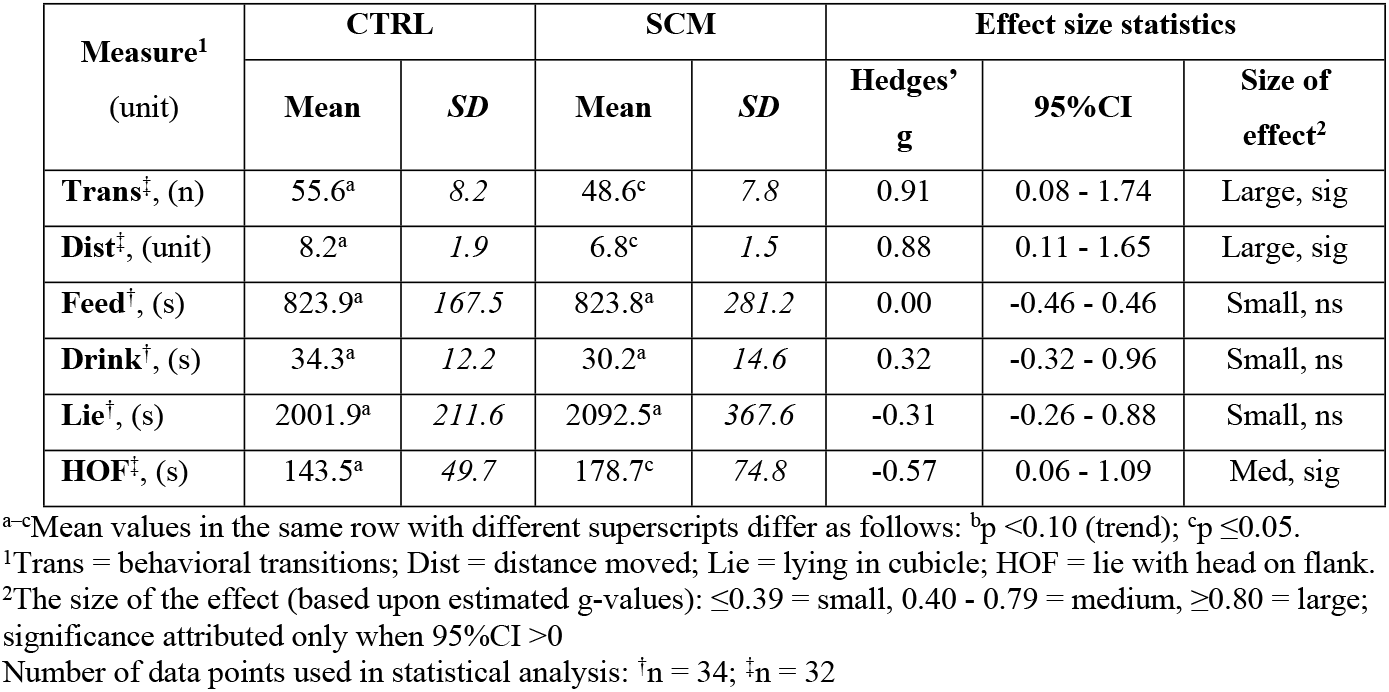
Measures of ‘core’ maintenance behavior recorded from pair-matched cows with (SCM) or without (CTRL) sub-clinical mastitis over 24h: analysed using paired t-tests. All units transformed to ‘per hour visible’.

| Measure<sup>1</sup><br>(unit) | CTRL |  | SCM |  | Effect size statistics |  |  |
| --- | --- | --- | --- | --- | --- | --- | --- |
|  | Mean | SD | Mean | SD | Hedges' g | 95%CI | Size of effect<sup>2</sup> |
| **Trans**<sup>‡</sup>, (n) | 55.6<sup>a</sup> | 8.2 | 48.6<sup>c</sup> | 7.8 | 0.91 | 0.08 - 1.74 | Large, sig |
| **Dist**<sup>‡</sup>, (unit) | 8.2<sup>a</sup> | 1.9 | 6.8<sup>c</sup> | 1.5 | 0.88 | 0.11 - 1.65 | Large, sig |
| **Feed**<sup>†</sup>, (s) | 823.9<sup>a</sup> | 167.5 | 823.8<sup>a</sup> | 281.2 | 0.00 | -0.46 - 0.46 | Small, ns |
| **Drink**<sup>†</sup>, (s) | 34.3<sup>a</sup> | 12.2 | 30.2<sup>a</sup> | 14.6 | 0.32 | -0.32 - 0.96 | Small, ns |
| **Lie**<sup>†</sup>, (s) | 2001.9<sup>a</sup> | 211.6 | 2092.5<sup>a</sup> | 367.6 | -0.31 | -0.26 - 0.88 | Small, ns |
| **HOF**<sup>‡</sup>, (s) | 143.5<sup>a</sup> | 49.7 | 178.7<sup>c</sup> | 74.8 | -0.57 | 0.06 - 1.09 | Med, sig |

<sup>a-c</sup>Mean values in the same row with different superscripts differ as follows: <sup>b</sup>p < 0.10 (trend); <sup>c</sup>p ≤ 0.05.
<sup>1</sup>Trans = behavioral transitions; Dist = distance moved; Lie = lying in cubicle; HOF = lie with head on flank.
<sup>2</sup>The size of the effect (based upon estimated g-values): ≤ 0.39 = small, 0.40 - 0.79 = medium, ≥ 0.80 = large; significance attributed only when 95%CI > 0
Number of data points used in statistical analysis: <sup>†</sup>n = 34; <sup>‡</sup>n = 32

**Table 6:** Differences in ‘core’ maintenance behavior identified between cows with (SCM) or without (CTRL) sub-clinical mastitis during 60 mins following morning milking (1hPostM1): parametric data (P) analysed using paired t-test, non-parametric data (NP) analysed using Wilcoxon SR test.

| Measure<sup>1</sup><br>(unit) | Unit | Control |  | SCM |  | Effect size statistics |  |  |
| --- | --- | --- | --- | --- | --- | --- | --- | --- |
|  |  | Mean | SD | Mean | SD | Hedge's g (bootstrap) | 95%CI | Size of effect<sup>2</sup> |
| **Trans**, (n) | NP | 99.1<sup>a</sup> | 34.8 | 71.7<sup>c</sup> | 24.6 | -0.89 | 0.21 - 1.62 | Large, sig |
| **Dist**, (unit) | P | 15.9<sup>a</sup> | 6.6 | 12.5<sup>b</sup> | 4.0 | -0.63 | -0.08 - 1.34 | Med, ns |

<sup>a-c</sup>Mean values in the same row with different superscripts differ as follows: <sup>b</sup>p < 0.10 (trend); <sup>c</sup>p ≤ 0.05.
<sup>1</sup>Trans = behavioral transitions; Dist = distance moved.
<sup>2</sup>The size of the effect (based upon estimated g-values): ≤ 0.39 = small, 0.40 - 0.79 = medium, ≥ 0.80 = large; significance attributed only when 95%CI > 0

##### (b) Physical activity

During both the 24h period, and 60 minutes following morning milking, the SCM cows were significantly less active than healthy controls; SCM cows performed fewer behavioral transitions (‘**Trans**’) and moved over a smaller distance (‘**Dist**’) (Tables 5 and 6).

### 3.2 Correlations Between Physiology and Behavior

#### 3.2.1 Assay validation

Parallelism (F_1,9_ = 3.46, p >0.05) was confirmed between serial dilutions of saliva (range: 1:4 to 1:64) and SAA standards (range: 0, 9.38, 18.75, 37.5, 75, 150, 300 ng/ml), indicating that the ELISA kit was suitable for use with bovine saliva. Recovery of 300 ng/ml SAA from a spiked saliva sample was 93.76 ± 4.63% (n = 10). The intra-assay CV was 3.09% (250.87 ± 7.75 ng/ml, n = 10) for QC_low_ and 4.68% (1360.33 ± 63.70 ng/ml, n = 10) for QC_high_. The inter-assay CV was 2.77% (246.06 ± 6.81 ng/ml, n = 2) for QC_low_ and 3.89% (1323.96 ± 51.43 ng/ml, n = 2) for QC_high_.

#### 3.2.2 SAA and SCC

The average SCC per group was: CTRL = 48.29 ± 28.33 (x1000 cells/ml); SCM = 351.12 ± 176.73 (x1000 cells/ml). A trend was found towards a significantly higher concentration of salivary SAA in the SCM cows (CTRL = 343.42 ± 269.60 ng/ml, SCM = 519.59 ± 315.43 ng/ml; t_1,12_ = 1.93, p = 0.076). A weak positive relationship was evident between SCC and salivary SAA levels (F_1,29_ = 8.81, p = 0.006, Figure 2).

**Figure 2:**
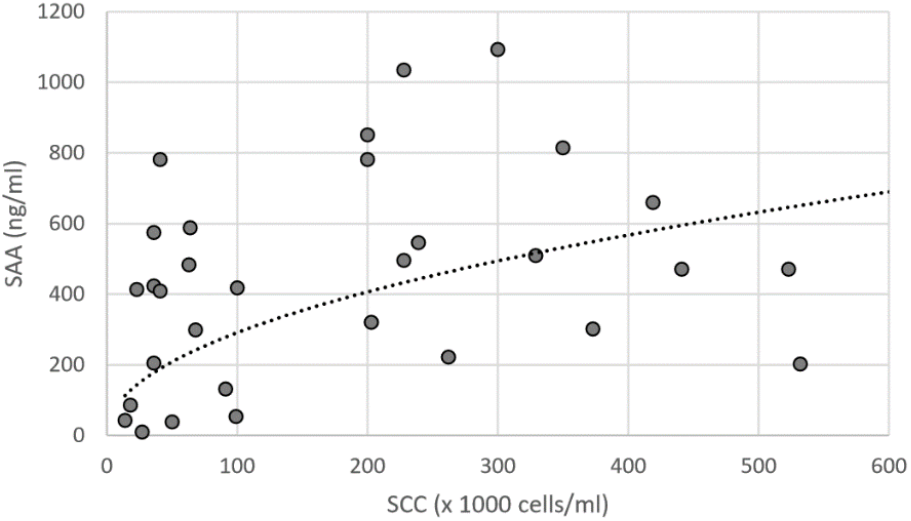
The significant positive relationship between somatic cell count (SCC) in milk and salivary serum amylase-A (SAA) in dairy cattle (R^2^ = 0.233; y = 31.715x^0.4815^)

#### 3.3.3. SAA, SCC and behavior

All behavioral measures significantly correlated with SAA and/or SCC are presented in Table 8. The majority of these relationships are weak as indicated by R^2^ values.

**Table 8:** Significant correlations (p<0.05; plus trends towards a correlation at p<0.10) between behavioral measures (24h data) and two markers of inflammation and mastitis infection, one from saliva (serum amyloid-A, SAA), and one from milk (somatic cell counts, SCC); given are curve estimation regression statistics (ANOVA, coefficient of determination) and the equation for the relationship (based upon the line of best fit).

| Behavioral measure<sup>1</sup><br>(unit) | Correlation with salivary SAA and SCC |
| --- | --- |
| **SocG** (n) | SAA: $F_{(1,29)} = 6.42$, $p = 0.017$; $R^2 = 0.181$; $y = 6.0091e^{-7E-04x}$ |
| **SocR** (n) | SAA: $F_{(2,28)} = 7.40$, $p = 0.003$; $R^2 = 0.346$; $y = 9E-06x^2 - 0.0061x + 5.057$<br>SCC: $F_{(2,31)} = 4.90$, $p = 0.014$; $R^2 = 0.240$; $y = 2E-05x^2 - 0.0081x + 5.259$ |
| **FocSocG** (n) | SAA: $F_{(1,29)} = 3.07$, $p = 0.090$; $R^2 = 0.096$; $y = 4.2715e^{-6E-04x}$ |
| **FocSocR** (n) | SAA: $F_{(2,28)} = 4.23$, $p = 0.025$; $R^2 = 0.232$; $y = 6E-06x^2 - 0.004x + 3.817$ |
| **CR** (n) | SAA: $F_{(1,29)} = 3.93$, $p = 0.057$; $R^2 = 0.119$; $y = 0.0003x + 0.165$ |
| **HPR** (n) | SCC: $F_{(1,31)} = 5.87$, $p = 0.021$; $R^2 = 0.159$; $y = -1.990x + 5.520$ |
| **HSG** (n) | SAA: $F_{(1,29)} = 6.89$, $p = 0.014$; $R^2 = 0.192$; $y = 2.3287e^{-9E-04x}$ |
| **HSR** (n) | SCC: $F_{(2,30)} = 3.14$, $p = 0.058$; $R^2 = 0.173$; $y = 6E-06x^2 - 0.0018x + 1.621$ |
| **%FocSocR+D** (%) | SAA: $F_{(1,29)} = 4.63$, $p = 0.040$; $R^2 = 0.138$; $y = -0.0155x + 52.078$<br>SCC: $F_{(1,32)} = 10.46$, $p = 0.003$; $R^2 = 0.246$; $y = 49.136e^{-7E-04x}$ |
| **%HBR+D** (%) | SAA: $F_{(1,29)} = 5.64$, $p = 0.024$; $R^2 = 0.163$; $y = 72.616e^{-5E-04x}$<br>SCC: $F_{(1,31)} = 8.81$, $p = 0.006$; $R^2 = 0.221$; $y = -7.36\ln(x) + 95.594$ |
| **%HPR+D** (%) | SAA: $F_{(2,22)} = 4.14$, $p = 0.030$; $R^2 = 0.227$; $y = -0.0002x^2 + 0.1952x - 0.092$ |
| **%HSR+D** (%) | SAA: $F_{(1,29)} = 3.291$, $p = 0.080$; $R^2 = 0.102$; $y = -0.015x + 34.655$ |
| **ExpSoc** (s) | SAA: $F_{(1,29)} = 4.34$, $p = 0.046$; $R^2 = 0.130$; $y = 0.0167x + 8.517$<br>SCC: $F_{(1,31)} = 5.58$, $p = 0.025$; $R^2 = 0.153$; $y = 15.532e^{-0.002x}$ |
| **%FB_Open** (%) | SAA: $F_{(1,28)} = 4.08$, $p = 0.053$; $R^2 = 0.127$; $y = 3.244\ln(x) + 67.752$<br>SCC: $F_{(1,32)} = 5.98$, $p = 0.020$; $R^2 = 0.158$; $y = 81.19e^{0.0003x}$ |
| **ExpEnv** (s) | SAA: $F_{(1,29)} = 5.96$, $p = 0.021$; $R^2 = 0.170$; $y = 8.690\ln(x) + 7.484$ |
| **Trans** (n) | SCC: $F_{(2,30)} = 5.50$, $p = 0.009$; $R^2 = 0.269$; $y = 0.002x^2 - 0.0934x + 59.272$ |
| **Dist** (unit) | SCC: $F_{(2,30)} = 4.11$, $p = 0.026$; $R^2 = 0.215$; $y = 4E-05x^2 - 0.021x + 9.146$ |
| **Feed** (s) | SAA: $F_{(1,29)} = 14.63$, $p = 0.001$; $R^2 = 0.335$; $y = 1023.5e^{-6E-04x}$ |
| **Drink** (s) | SAA: $F_{(1,29)} = 22.18$, $p < 0.001$; $R^2 = 0.433$; $y = -8.379\ln(x) + 79.816$ |
| **Lie** (s) | SAA: $F_{(1,29)} = 9.10$, $p = 0.005$; $R^2 = 0.239$; $y = 0.5032x + 1848.3$<br>SCC: $F_{(2,31)} = 2.637$, $p = 0.088$; $R^2 = 0.145$; $y = -0.0019x^2 + 1.1955x + 1956.8$ |

<sup>1</sup>SocG = all social behavior given; SocR = all social behavior received; FocSocG = agonistic behavior given; FocSocR = agonistic behavior received; CR = non-contact challenge receive; HPR = head push receive; HSG = head swipe give; HSR = head swipe receive; %FocSocR+D = percentage of occasions when receipt of agonism lead to displacement; %HBR+D = percentage of occasions when head butt receipt lead to displacement; %HPR+D = percentage of occasions when head push receipt lead to displacement; %HSR+D = percentage of occasions when head swipe receipt lead to displacement; ExpSoc = explore social; %FB\_Open = percentage of time at feed barrier: open rail; ExpEnv = explore environment; Trans = behavioral transitions; Dist = distance moved.

##### (a) Total social interactions and agonistic interactions

Significant positive relationships were found between salivary SAA and total social interactions received (**‘SocR’**) and total agonistic interactions received (**‘FocSocR’**), indicating that cows with higher levels of salivary SAA cows received more social interactions in total, and also more agonistic interactions. Significant negative correlations were found between salivary SAA and total social interactions given (**‘SocG’)** and head swipes given (**‘HSG’**), with a trend towards significance for total agonistic interactions given (**‘FocSocG’**). This indicates that cows with higher levels of salivary SAA cows initiated fewer social interactions in total, including fewer agonistic interactions.

The significant relationship between SCC and total social interactions received (**‘SocR’**) was best modelled using a quadratic function. This described a negative association at low SCC levels and a positive association at higher levels, such that as cell counts rose in cows with low SCC (i.e. healthy CTRL cows) the tendency to receive social interactions would decrease, but in those cows with higher SCC (i.e. SCM cows) as cell counts rose further the tendency to receive social interactions increased. The significant positive correlation between SCC and the receipt of head swipes (‘**HSR’)** indicates that cows with higher SCC received more agonistic head swipes.

##### (b) Social reactivity

Significant negative relationships were found between measures of social reactivity, (i.e. the likelihood that a cow would be displaced/move away after receiving an agonistic interaction), and both inflammatory markers. This is indicative that cows with higher levels of salivary SAA and/or SCC demonstrate a reduced tendency to be displaced following the receipt of social stimuli, including total agonistic interactions (‘**%FocSocR+D** ‘), and specifically to head butts (‘**%HBR+D’)** (SAA and SCC), and head swipes received (‘%HSR+D’) (SAA).

##### (c) Social and environmental exploration, and feed barrier preference

Salivary SAA was significantly positively associated with both social and environmental exploration, such that cows with higher SAA levels performed more exploratory behavior. SCC levels, however, were significantly negatively associated with social exploration, such that cows with higher cell counts explored other cows less. Salivary SAA and SCC were significantly positively correlated with proportion of time spent at the open-rail feed barrier (**‘%FB_Open’**), indicating that with rising levels of either inflammation measure, cows became more likely to feed at the open-rail feed barrier rather than the self-locking barrier.

##### (d) Physical activity, feeding, drinking, and lying

The significant relationships between SCC and both the number of behavioral transitions and distance covered were best modelled using quadratic functions. These describe an initial decline in both measures of physical activity as SCC increased to approximately 300 (x1000 cells/ml), followed by an increase in the measures as SCC levels continued to rise (Figure 3).

**Figure 3:**
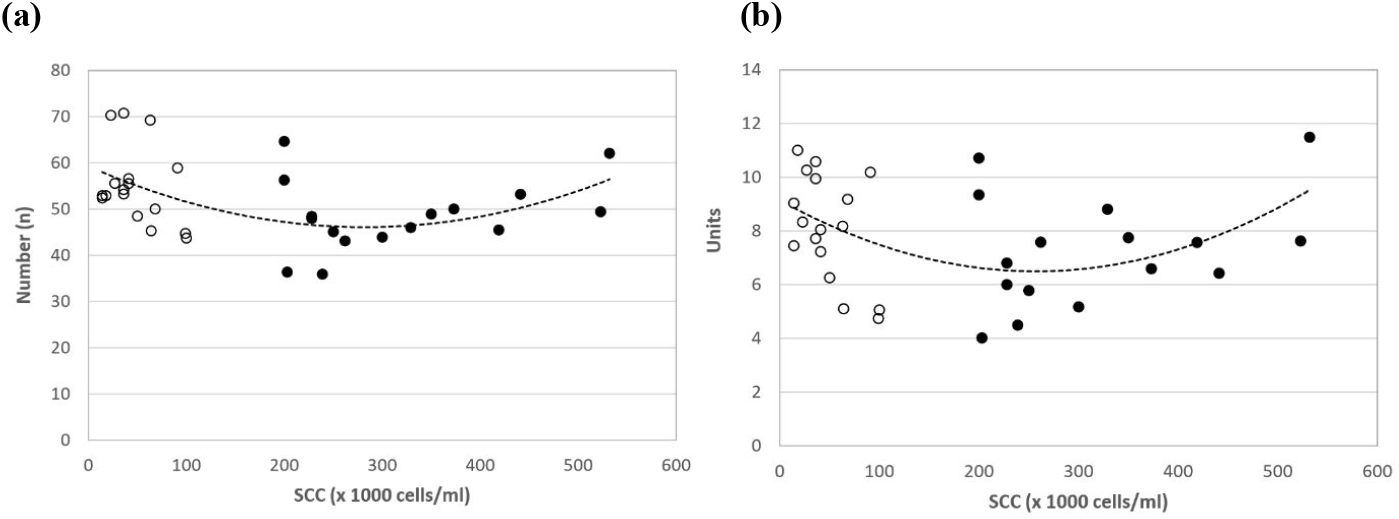
Quadratic relationship between somatic cell count (SCC) and two measures of activity in cows with (SCM = black), and without (CTRL = open circle) subclinical mastitis: (a) behavioral transitions (‘Trans’); (b) distance covered (‘Dist’).

Salivary SAA was significantly negatively correlated with feeding and drinking, and significantly positively correlated with time spent lying; this indicates that, within our study cohort, that as SAA levels increased the cows spent less time feeding and drinking, and more time lying.

## 4. DISCUSSION

The purpose of this study was to identify potential changes in salivary serum amyloid-A (SAA) and behavior in cows with spontaneously occurring subclinical mastitis as compared to pair-matched, healthy controls. Salivary SAA was found to be higher in individuals with subclinical mastitis, suggesting that it may have potential as a marker of low-level systemic inflammation in dairy cows. Higher salivary SAA levels were also correlated with several predicted behavioral changes. Furthermore, cows with subclinical mastitis displayed a reduction in the performance and receipt of various social behaviours (here categorised as ‘luxury’ behaviors), lower ‘social reactivity’ (i.e. they were less likely to be displaced following the receipt of agonism), a reduction in physical activity, but no difference in feeding, drinking or lying duration (‘core maintenance’ behaviors).

### 4.1. Changes in salivary SAA

We report here, for the first time, a positive (although weak) correlation between SCC and salivary SAA. Positive associations between SCC and non-salivary SAA have previously been reported from cows with clinical and sub-clinical mastitis (serum: de Boyer Des Roches et al., 2017; milk: O’Mahony et al., 2006; Akerstedt et al., 2007; Pyörälä et al., 2011). SAA in saliva thus offers potential as a non-invasive means of detecting subclinical mastitis, but further studies will be required to substantiate our preliminary findings. Different bacterial strains, for example, can cause mastitis of different duration and degree (Verbeke et al., 2014), and it is possible that the concentration of SAA in saliva will vary accordingly. A case in point is Pyörälä et al.’s (2011) finding of significant differences in SAA (milk) collected from cows with spontaneous mastitis caused by different pathogens: low SAA was associated with *A. pyogenes*, while high concentrations were associated with *E. coli*. Consequently, further investigations will need to establish how, and when, salivary SAA levels change over the time course of an infection, with different pathogens, to substantiate its diagnostic potential as an inflammatory marker for mastitis and other diseases.

### 4.2. Changes in Luxury Behaviors and correlations with salivary SAA

‘Luxury’ behaviors, as used here, are those deemed non-essential for survival within the short-term and, as such, we predicted that during a subclinical infection they would be down-regulated. Within this category we included both social and non-social behaviors.

#### 4.2.1. Social interactions

Over the 24h period, total social behavior, given or received, remained unaltered between the two groups. Subclinically mastitic cows did, however, receive fewer non-contact challenges and more head swipes. Head swipes are most commonly observed at the feed barrier as a means of displacing competitors, and our result is in agreement with previous findings. Sick cows are typically reported to be displaced more frequently than healthy cows when feeding, as would result from head swipes or other agonistic behavior (Schirmann et al., 2016; Lomb et al., 2018; Neave et al., 2018;).

In the 60 minutes following morning milking, that is during the period of peak feeding activity in the study herd, other differences emerged; cows with subclinical mastitis received fewer social interactions in total, and specifically fewer head butts and body pushes, than the healthy controls. No corresponding reduction in the receipt of agonistic interactions involving physical contact were observed over the 24h period. It is therefore possible that the subclinically mastitic cows were avoiding peak feeding when aggression and competition for feed are highest, as do low-ranking cows (Val-Laillet et al. 2008).

Interestingly, we also found a positive correlation between salivary SAA and total social interactions, and agonistic interactions, received, indicating that those individuals with higher levels of systemic inflammation received more agonistic interactions. However, we cannot be sure whether SAA upregulation occurred in the cows with subclinical mastitis due to infection or following exposure to social stress. Upregulation of C-Reactive Protein (another acute phase protein known to increase during illness and stress) has, for example, been reported in zoo-housed gorillas following an aggressive encounter (Fuller & Allard, 2018). In the current study, saliva samples were collected the day after behavioral video recordings. Therefore, an elevation in SAA could reflect agonistic encounters experienced during the previous day, independent of health status. Further study should address this potential confound.

In line with the prediction that social behaviors should decrease with subclinical mastitis, affected cows performed fewer head butts than healthy cows, and SAA was negatively (albeit loosely) correlated with the delivery of total social and agonistic interactions, including head swipes in particular. Our results thus provide additional behavioral detail in support of existing literature which reports that cows with clinical (and subclinical) conditions often perform fewer agonistic interactions and competitive displacements from the feed-bunk (Galindo & Broom, 2002; Huzzey et al., 2007; Patbandba et al., 2012; Sepúlveda-Varas et al., 2014; 2016), and from cubicles (Jensen & Proudfoot, 2017), than healthy individuals.

The relative social ranks of study cows and their interaction partners were not determined in the current study. It is possible, and an interesting focus for further study, that the relative social rank of an individual could be influenced by the effects of disease due to a loss of competitive vigour and motivation. High-ranking animals characteristically displace lower rankers cows from the feed barrier (DeVries et al., 2004; Huzzey et al., 2006), and lower rankers adjust their feeding patterns accordingly (DeVries et al., 2004).

#### 4.2.2. Social reactivity

We found that over the 24h period, cows with subclinical mastitis were less likely to move away/be displaced than matched healthy cows following the receipt of agonism, labelled here as less ‘socially reactive’. Correspondingly, reactivity declined with increasing inflammation levels as demonstrated by a negative correlation between SAA levels and percentage displacement following the receipt of agonistic interactions overall, and specifically following the receipt of head butts and head swipes. Similarly, SCC was negatively correlated with the percentage displacement following receipt of agonistic interactions overall, and specifically of head butts, such that cows with higher SCC were less likely to more away after being head butted, and/or following receipt of all agonistic interactions, than cows with lower counts. These results appear, at first hand, to contradict previous findings of sick cows being less motivated to compete for access to feed based on displacement (e.g. Huzzey et al, 2006; 2007; Goldhawk et al., 2009); however, where a cow positions herself at the feeder may, in turn, be influenced by the social rank of others already feeding, as reported by Manson & Appleby (1990). They found that cows of the most dissimilar rank (low/high) maintained a greater distance from each other than cows of similar rank. It is thus conceivable that subclinically sick cows proactively avoided feeding positions next to individuals of higher competitive vigour, and preferentially selected the company of other sick or lower ranking cows. Subsequently, the receipt of moderate competitive aggression from an individual, with similarly moderate motivation to compete, may have been tolerable and thus not provoked displacement. This is beyond the reach of the current data as we did not determine the relative social ranks of our focal cows and their interaction partners, but warrants further investigation. As others before us, we can therefore not rule out that this and our other findings on social behavioral differences reflect underlying differences in social status that have led to differences in health status. We here provide a first report of a reduction in social reactivity in cows with subclinical mastitis. By matching cows by parity we sought some control over potentially confounding social rank biases. Further study is needed to substantiate differences in social interactions as markers of subclinical mastitis and other diseases.

#### 4.2.3. Social avoidance and proximity

In the current study, cows with subclinical mastitis performed less social exploration than healthy controls during both the 24h period and the focussed 60 min following morning milking, which indicates social avoidance or withdrawal. Furthermore, we observed a negative correlation between this measure and SCC, such that cows with higher counts spent less time exploring (sniffing) others. Sickness-driven social avoidance is well documented in lab-animals and humans and is predicted via the action of pro-inflammatory cytokines on the central nervous system (Kent et al., 1992; Bluthe et al., 1996; Dantzer & Kelley, 2007; Arakawa et al., 2010). Due to the limited opportunity for social avoidance to occur within intensive systems it has been relatively understudied in farm animals. We also found an unexpected, if weak, positive correlation between SAA and social exploration, which may, at least partially, be explained by the presence of both pre-clinical and post-clinical individuals amongst our subclinical group. De Boyer Des Roches et al. (2017; 2018) report behavioral changes including reduced environmental attentiveness during the pre-clinical phase of an experimentally-induced mastitis model (i.e. prior to SCC and serum SAA upregulation), and during the acute phase (coinciding with raised levels of SCC and SAA), but not during the post-clinical remission phase (when high levels of SCC and SAA are evident). This suggests that serum SAA may peak during post-clinical remission, rather than during the acute phase of inflammation. Further study is therefore required to investigate when, and how, social exploration changes throughout the course of spontaneously-occurring mastitis, and other diseases, to confirm its usefulness in the detection of subclinical disease states.

We found that over the 24h period, cows with subclinical mastitis spent a greater proportion of their time feeding at the open-rail section of the feed barrier than healthy controls. A weak positive correlation was also found between this measure and SAA, which suggests that this preference increases with systemic inflammation. Barrier design is known to influence the incidence of agonistic behavior relating to feed access. Self-locking yokes have vertical bars which separate the necks of adjacent cows, and these are better at reducing competitive interactions (usually measured by displacements) compared to open post-and-rail barriers (Endres et al., 2005; Huzzey et al., 2006). Our results therefore appear to contradict previous findings. However, other factors are likely to also contribute to the choice of feeding location. In the open section cows have better visibility and are more quickly able to withdraw from potential agonistic interactions. Our finding that subclinically mastitic cows spent a greater proportion of their time than the healthy cows at the feed barrier in close contact with two (flanking) herdmates may be at odds with the expectation of sickness-driven social avoidance. However, sick cows may also feel more vulnerable and therefore seek to feed in company. Future longitudinal studies could determine whether individuals change their feeding behavior in relation to health status, including whether they opt to feed amongst others of lower rank when their own health is compromised.

#### 4.2.4. Allogrooming

In agreement with our predictions relating to the basis of behavioral priorities during infection, cows with subclinical mastitis performed less allogrooming than the healthy cows over the 24h period, and during the 60 min following morning milking they themselves were also allogroomed less. Previous studies have reported similar findings in relation to social rank, with low-ranking cows performing and receiving less allogrooming than high rankers (Napolitano et al., 2009), and allogrooming decreasing more in low-ranking cows under conditions of increased competition (Val-Laillet et al., 2008). For health-related changes, Galindo & Broom (2002) found lame cows to be allogroomed more than non-lame cows, which is contrary to our result in subclinically mastitic cows. Their finding was interpreted as a self-instigated coping strategy triggered by pain/discomfort associated with clinical lameness, which arguably applies less to subclinical mastitis.

Since social grooming serves a variety of functions in cattle, including roles in hygiene, pleasure, maintenance of social bonds and lowering social tension (Sato et al., 1991; 1993), it is possible that prolonged suppression of allo-grooming, for example during a low-level chronic inflammatory condition, could negatively affect cow welfare and health itself.

#### 4.2.5. Non-social behavior

##### (a) Self-grooming

Against expectation, we did not find a difference in brush use between cows with and without subclinical mastitis, nor a correlation between this behavior and SAA. Although grooming itself is a comfort activity that healthy cows are highly motivated to perform (McConnachie et al., 2018), brush use has been shown to decrease during non-mastitic disease (Toaff Rosenstein et al., 2016; Mandel et al., 2017; Weigele et al., 2017). The brush in our study was located central to many resources including the feed barrier, a water trough and cubicles and was, therefore, readily accessible to all cows with minimal effort. In addition to our relatively small sample size, a trade-off between brush location and the sensitivity of brush use for detecting stress and morbidity (Mandel et al., 2013; 2017) may help to explain why no decline was detected in the current study.

We observed cows with subclinical mastitis to perform more self-grooming (including self-licking) than controls immediately following morning milking, presumably as a response to mild udder discomfort or as a substitute for allo-grooming (see section 4.2.4). However, this observation is at odds with existing literature that reports self-licking to remain unchanged (Siivonen et al., 2011), or even decrease (Fogsgaard et al., 2012), during clinical mastitis. Based upon our results the potential for the use of self-grooming as a marker of subclinical mastitis remains inconclusive and requires further investigation.

##### (b) Environmental exploration

The weak positive correlation between SAA and environmental exploration described in the current study was unexpected since exploratory behavior was predicted to decline with increasing inflammation. As our study did not use an experimental model of induced mastitis we do not know whether our focal cows comprised a mixture of early-stage, pre-clinical mastitic cows (potentially associated with low SAA and sickness-driven reductions in environmental exploration), and individuals in post-clinical remission (potentially associated with high SAA and normal baseline values) (as discussed in Section 4.2.3). Ruminants generally display low baseline levels of exploratory behavior when maintained in intensive housing, since it is a largely unstimulating environment (De Rosa et al., 2009). Against expectation, we also did not find a reduction in environmental exploration in our subclinically mastitic cows over the 24 hours. They did, however, explore less than controls during the 60 min following morning milking. Environmental exploration, as defined in our study, included sniffing the feed as well as pen fittings and cubicle sand. Lower levels of exploration in cows with subclinical mastitis after morning milking may therefore also reflect avoidance of feeding at peak time.

### 4.3. Core Maintenance Behavior

#### 4.3.1. Activity

Our finding that cows with subclinical mastitis made fewer behavioral transitions and moved over a shorter distance than healthy cows is in agreement with our predictions, and with other studies that describe reduced activity prior to the clinical diagnosis of mastitis (Kester et al., 2015; Stangaferro et al., 2016; Veissier et al., 2017; King et al., 2018). The quadratic relationships between SCC and both ‘behavioral transitions’ and ‘distance moved’ described in the current study are of interest because mastitic cows have also been reported to display increased activity (Siivonen et al., 2011; Medrano-Galarza et al., 2012), presumably due to udder discomfort and an associated reduction in lying time. Jadhav et al. (2018) argue that the threshold SCC value to delineate subclinical mastitis from normal should be 310, rather than 200 (x1000 cells/ml), as conventionally judged (e.g. Madouasse et al., 2010). This higher value closely corresponds with the parabola vertex in both quadratic plots (Figure 3), of approx. 300 (x1000 cells/ml); i.e. the point at which activity once again begins to rise.

#### 4.3.2. Feeding/Drinking

Changes in feeding behavior have long been used to diagnose the onset of illness (Weary et al., 2009). We found a negative correlation between SAA and feeding duration, as would be predicted with sickness. Although we observed a negative correlation between SAA and feeding duration, as would be hypothesised to occur with sickness, the average inflammatory response within our SCM group overall was not sufficiently pronounced to trigger obvious anorexia, as compared to CTRL cows. Sepúlveda-Varas et al. (2016) observed a decrease in feed intake (but not duration) prior to the diagnosis of clinical mastitis which may be attributed to underlying malaise. González et al. (2008) reported individual variability in feeding behavior relating to spontaneously occurring udder disorders. Some cows demonstrated a decrease in feeding duration with the onset of mastitis, while others showed no change. It is possible that aspects of feeding behavior, other than duration, may have been affected. Barn-housed cattle demonstrate highly synchronised feeding activity, with large peaks in both feeding and social competition coinciding with fresh food delivery, and smaller peaks following milking (DeVries & von Keyserlingk, 2005; Dollinger & Kaufmann, 2013). Mastitic cows, presumably to avoid adverse social interactions, have been shown to feed at less popular times such as early afternoon (Schirmann et al., 2016). While we found no direct effect on feeding duration, our findings on social interactions after morning milking, reported above, may be indicative of cows with subclinical mastitis avoiding peak feeding times.

Water and feed intake are positively related in cattle (Kume et al., 2010); however, drinking tends to be less affected by health than feeding (Hart, 1988). Water is more immediately vital for maintaining bodily functions (Kyriazakis and Tolkamp, 2011), and since drinking takes less time than food consumption it is less prone to being disrupted by social competition at the trough (Huzzey et al., 2007). Although a reduction in water consumption has been reported in cows with mastitis (Lukas et al., 2008; Siivonen et al., 2011), and we observed a moderate negative correlation between SAA and drinking duration, the level of systemic inflammation within our subclinically sick cows may have been too low, and/or our sample size too small, for a difference to emerge. Overall, the persistence of feeding and drinking levels in our subclnically mastitic cows fits the prediction that core maintenance behavior is conserved during early mastitic disease.

#### 4.3.3. Lying

No difference in lying duration was found between our two groups. Lying is a highly prioritised behavior in cattle due to its importance in rumination (Jensen et al., 2004; 2005; Munksgaard et al., 2005) and dairy cows spend approximately 11h/day recumbent (Ito et al., 2009; 2010). Increased lying duration, as a means of conserving energy and facilitating recovery, is a key adaptation for sickness, and a positive correlation between SAA and lying was found in our cows. Although extended lying duration has been frequently reported during non-mastitic clinic conditions (Toaff Rosenstein et al., 2016; Weigele et al., 2017; Barragan et al., 2018), lying may actually decrease during mastitis (Yeiser et al., 2012; Fogsgaard et al., 2012; 2015; Medrano-Galarza et al. 2012;), most likely due to udder pain (Cyples et al., 2012).

We also observed that cows with subclinical mastitis lay with their heads held against their flank more than the healthy controls. This posture is primarily associated with rapid eye movement (REM) sleep, but cows are also known to display non-rapid eye movement (NREM) sleep and drowsing in this position (Ternman et al., 2013), and NREM (deep) sleep often increases during infection (Bryant et al., 2004; Opp, 2005). Crucially, the action of pro-inflammatory cytokines during the sickness response also predicts postural changes, such as ‘curling up’, that reduce surface area and associated loss of body heat. Lying with neck and head bent back against and resting on the body achieves a reduction in surface area in cows, and as such fits with predictions for adaptive behavioral changes during sickness. On the basis of our study, lying duration itself appears less promising for the detection of sub-clinical mastitis than our novel finding of a difference in lying posture.

### 4.4 Limitations

This study formed a preliminary investigation, to identify behaviors with potential for use as a marker of sub-clinical mastitis, with the intention of informing a wider schedule of focused research; longitudinal studies are now required to track changes in key behaviors with deviations in the health status of individual cows. Due to an absence of pre-existing literature (luxury behavior, in this context, has been understudied), a large number of measures were recorded over 24h. Logistically this was time consuming and limited the amount of data available per cow; priority was given to sample size. The analysis of a single 24h period for each cow provided a snapshot of one day, It is therefore not possible to conclude that behaviors observed were typical of that cow at that level of SCC. Although we health-checked study cows for inclusion in the study, as described in the Methods, to rule out additional clinical pathologies we could not determine subclinical conditions other than mastitis, nor could we establish whether cows classified as having subclinical mastitis were in the pre-clinical phases of disease or in remission.

## 5. CONCLUSIONS

By studying in detail the behavior of healthy cows and cows with spontaneously occurring subclinical mastitis, pair matched by reproductive parameters and parity, over two time periods (24h and 60 min following morning milking) we found that subclinical mastitis was associated with reductions in activity, social exploration, the receipt of non-agonistic social behavior, social reactivity (probability of displacement following receipt of agonism), and an increase in the receipt of head swipes, compared to clinically healthy control cows. Many social interactions can be considered ‘luxury’ behaviors during sickness, and here we provide preliminary findings that suggest that several social measures change at subclinical levels of mastitis, as predicted, whereas ‘core’ maintenance activities (including feed, drink and lie) did not, and luxury behaviors therefore offer greater potential for use in early disease detection. The wider social context and, specifically the rank relationships between study cows and their interaction partners, however, were beyond the scope of this investigation, and further study is needed on the interactions between social rank, health status and behavior in cows.

Cows with subclinical mastitis displayed a different feeding pattern. They spent a greater proportion of their feeding time in direct contact with two neighbours, and a lower proportion of time feeding at the self-locking feed barriers, than the healthy cows. Although a positive relationship between SCC and salivary SAA was observed, and several correlations between SAA and behavioral measures were identified in a direction consistent with sickness behavior (including positive correlations with lying duration and the receipt of total agonism, and negative correlations with feeding, drinking, the performance of total social and agonistic behavior, and social reactivity), the majority of associations were relatively weak.

With this study, we have taken initial steps to identify physiological and detailed behavioral changes associated with subclinical mastitis; a number of our findings are consistent with predictions for low-level sickness responses. Social behavior is fundamentally dependent upon the wider social environment and necessitates interactions between focal animals and other individuals within the herd. We consider the preliminary identification of behavioral differences in a small group of cows within a complex and dynamic social environment very encouraging. We now recommend that observations be replicated in longitudinal studies, tracking individuals, and larger data sets to substantiate and refine our findings.

## ACKNOWLEDGEMENTS

This study was funded by The John Oldacre Foundation (grant code R12518-102). We would also like to thank the staff at Wyndhurst Farm for facilitating our research.

---

**The STROBE-Vet statement checklist.**
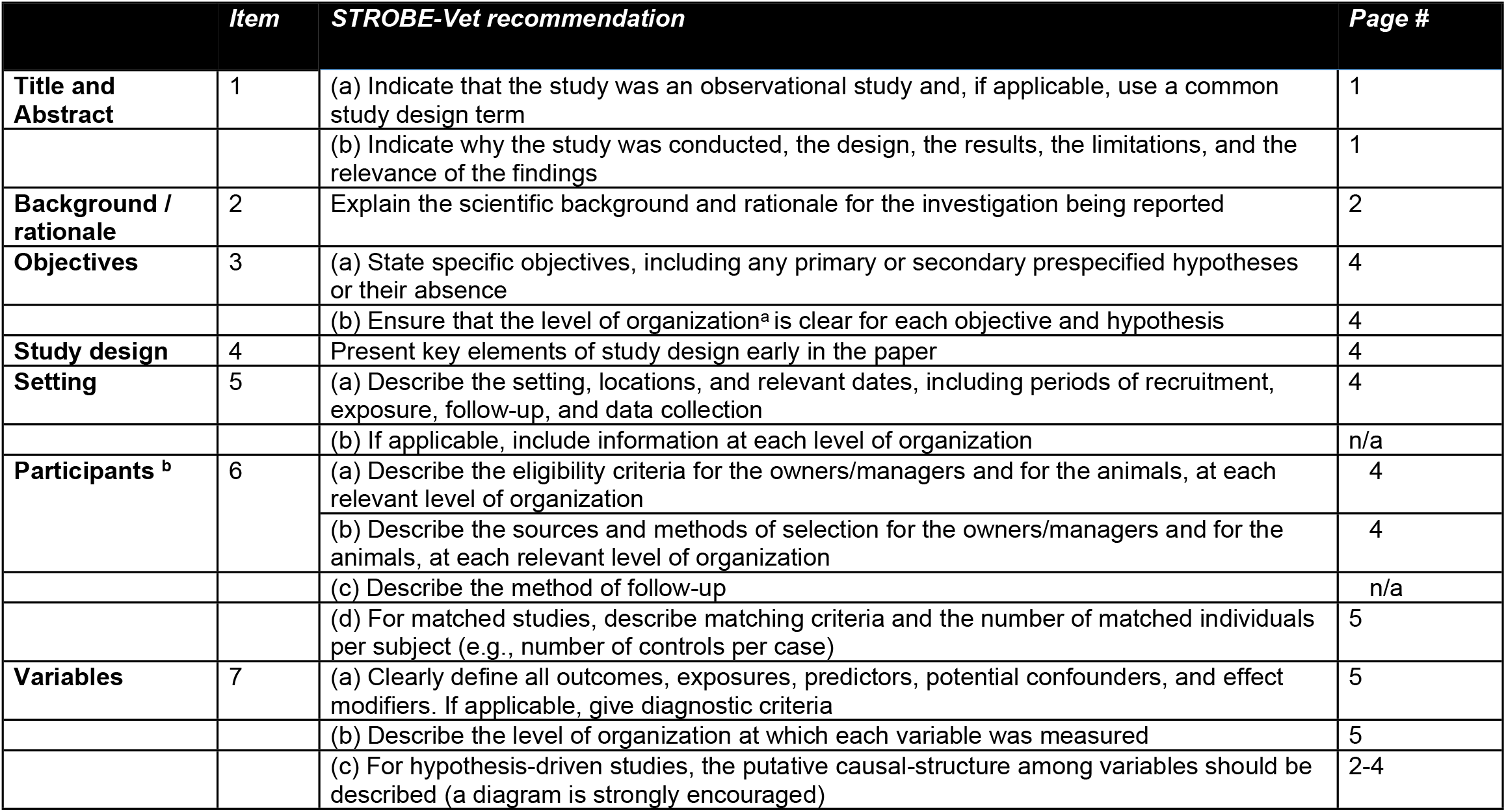
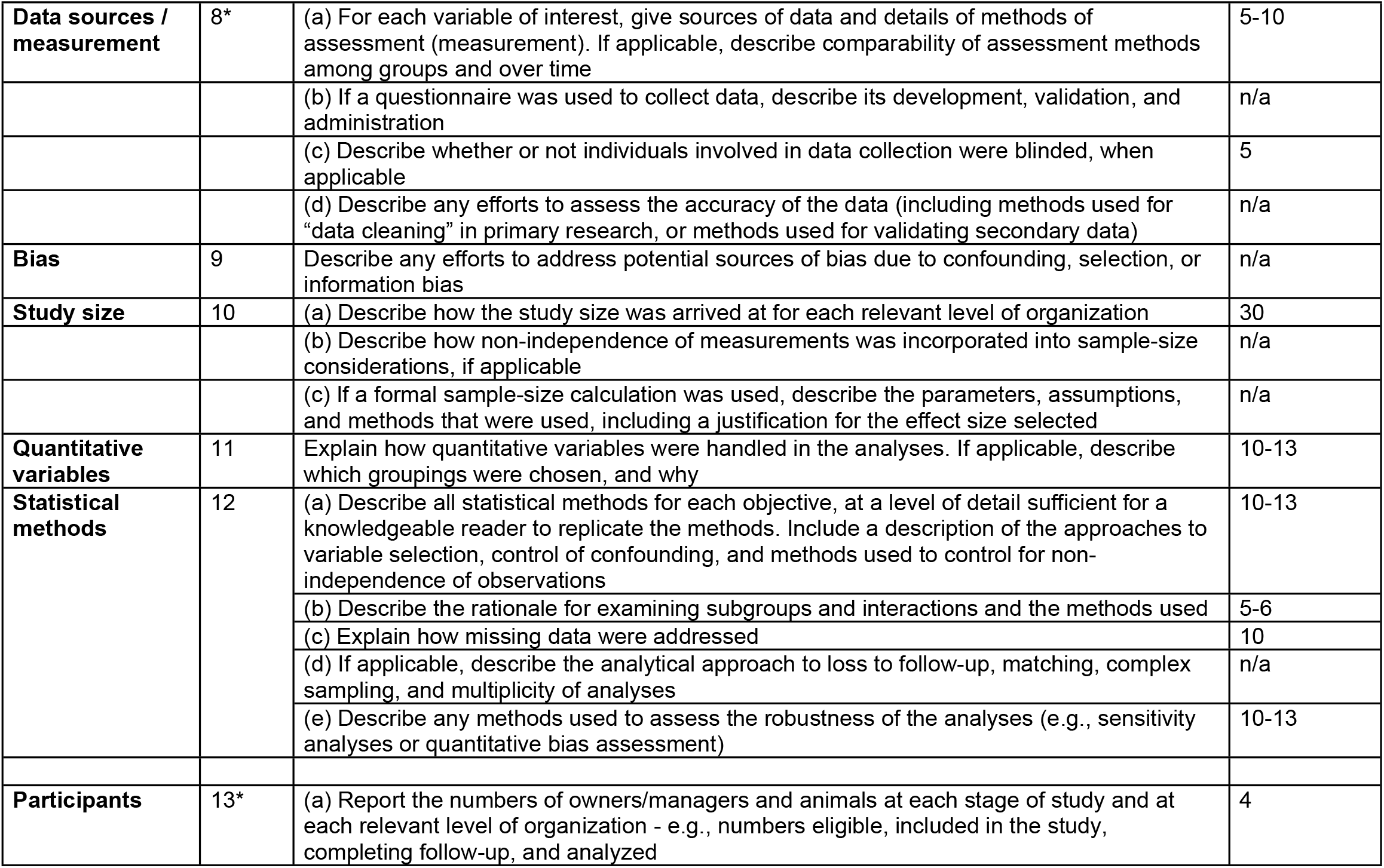
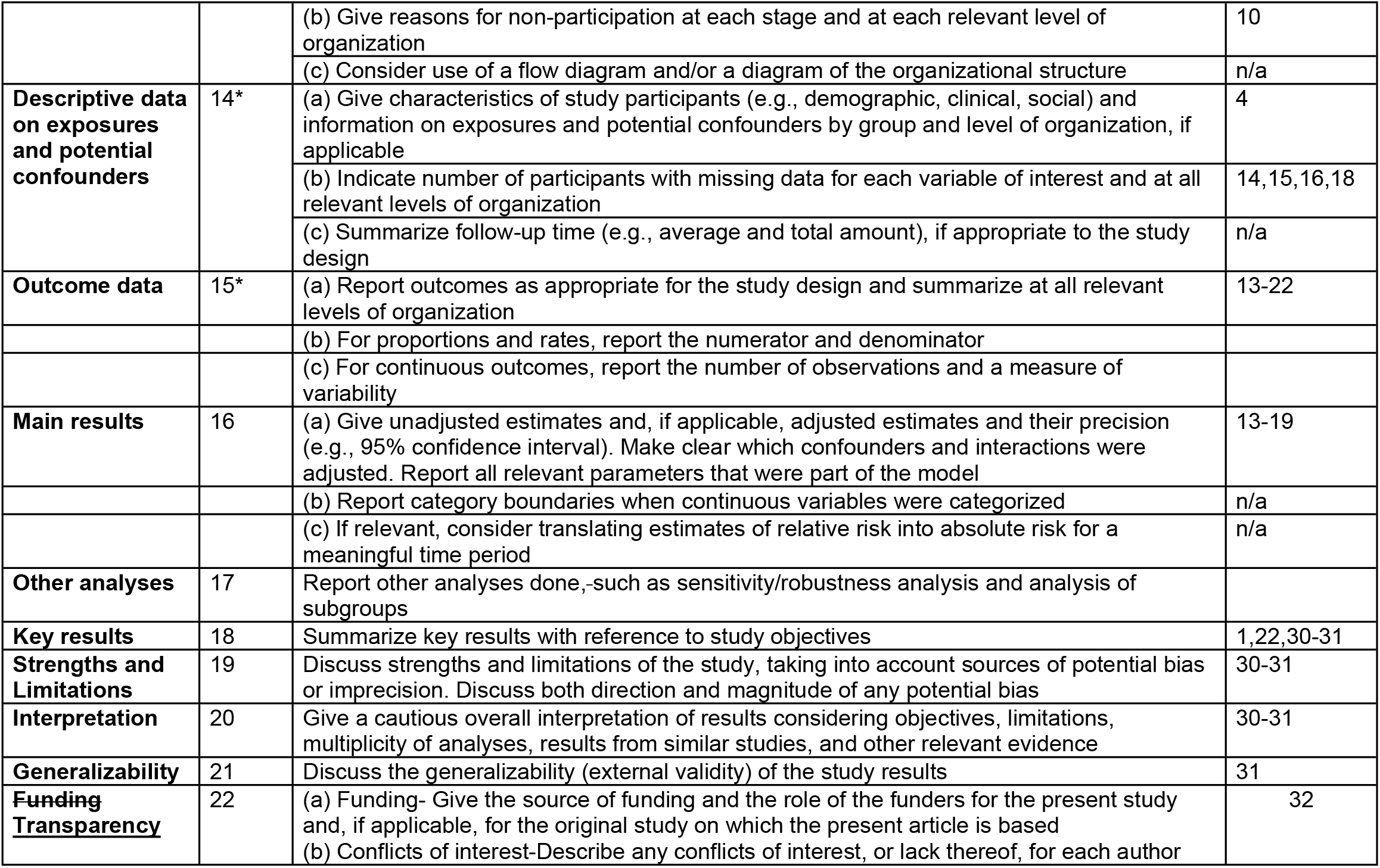
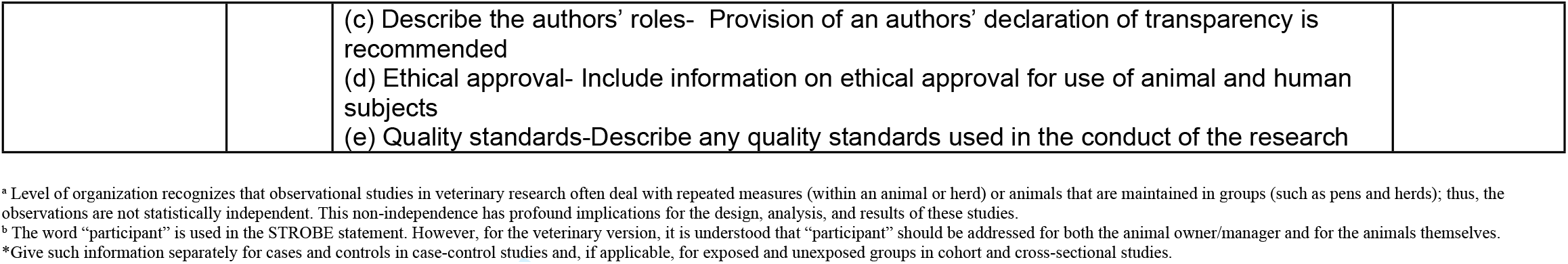

|  | Item | STROBE-Vet recommendation | Page # |
| --- | --- | --- | --- |
| Title and Abstract | 1 | (a) Indicate that the study was an observational study and, if applicable, use a common study design term | 1 |
|  |  | (b) Indicate why the study was conducted, the design, the results, the limitations, and the relevance of the findings | 1 |
| Background / rationale | 2 | Explain the scientific background and rationale for the investigation being reported | 2 |
| Objectives | 3 | (a) State specific objectives, including any primary or secondary prespecified hypotheses or their absence | 4 |
|  |  | (b) Ensure that the level of organization<sup>a</sup> is clear for each objective and hypothesis | 4 |
| Study design | 4 | Present key elements of study design early in the paper | 4 |
| Setting | 5 | (a) Describe the setting, locations, and relevant dates, including periods of recruitment, exposure, follow-up, and data collection | 4 |
|  |  | (b) If applicable, include information at each level of organization | n/a |
| Participants <sup>b</sup> | 6 | (a) Describe the eligibility criteria for the owners/managers and for the animals, at each relevant level of organization | 4 |
|  |  | (b) Describe the sources and methods of selection for the owners/managers and for the animals, at each relevant level of organization | 4 |
|  |  | (c) Describe the method of follow-up | n/a |
|  |  | (d) For matched studies, describe matching criteria and the number of matched individuals per subject (e.g., number of controls per case) | 5 |
| Variables | 7 | (a) Clearly define all outcomes, exposures, predictors, potential confounders, and effect modifiers. If applicable, give diagnostic criteria | 5 |
|  |  | (b) Describe the level of organization at which each variable was measured | 5 |
|  |  | (c) For hypothesis-driven studies, the putative causal-structure among variables should be described (a diagram is strongly encouraged) | 2-4 |
| **Data sources / measurement** | 8* | (a) For each variable of interest, give sources of data and details of methods of assessment (measurement). If applicable, describe comparability of assessment methods among groups and over time | 5-10 |
|  |  | (b) If a questionnaire was used to collect data, describe its development, validation, and administration | n/a |
|  |  | (c) Describe whether or not individuals involved in data collection were blinded, when applicable | 5 |
|  |  | (d) Describe any efforts to assess the accuracy of the data (including methods used for “data cleaning” in primary research, or methods used for validating secondary data) | n/a |
| **Bias** | 9 | Describe any efforts to address potential sources of bias due to confounding, selection, or information bias | n/a |
| **Study size** | 10 | (a) Describe how the study size was arrived at for each relevant level of organization | 30 |
|  |  | (b) Describe how non-independence of measurements was incorporated into sample-size considerations, if applicable | n/a |
|  |  | (c) If a formal sample-size calculation was used, describe the parameters, assumptions, and methods that were used, including a justification for the effect size selected | n/a |
| **Quantitative variables** | 11 | Explain how quantitative variables were handled in the analyses. If applicable, describe which groupings were chosen, and why | 10-13 |
| **Statistical methods** | 12 | (a) Describe all statistical methods for each objective, at a level of detail sufficient for a knowledgeable reader to replicate the methods. Include a description of the approaches to variable selection, control of confounding, and methods used to control for non-independence of observations | 10-13 |
|  |  | (b) Describe the rationale for examining subgroups and interactions and the methods used | 5-6 |
|  |  | (c) Explain how missing data were addressed | 10 |
|  |  | (d) If applicable, describe the analytical approach to loss to follow-up, matching, complex sampling, and multiplicity of analyses | n/a |
|  |  | (e) Describe any methods used to assess the robustness of the analyses (e.g., sensitivity analyses or quantitative bias assessment) | 10-13 |
| **Participants** | 13* | (a) Report the numbers of owners/managers and animals at each stage of study and at each relevant level of organization - e.g., numbers eligible, included in the study, completing follow-up, and analyzed | 4 |
|  |  | (b) Give reasons for non-participation at each stage and at each relevant level of organization | 10 |
|  |  | (c) Consider use of a flow diagram and/or a diagram of the organizational structure | n/a |
| **Descriptive data on exposures and potential confounders** | 14* | (a) Give characteristics of study participants (e.g., demographic, clinical, social) and information on exposures and potential confounders by group and level of organization, if applicable | 4 |
|  |  | (b) Indicate number of participants with missing data for each variable of interest and at all relevant levels of organization | 14,15,16,18 |
|  |  | (c) Summarize follow-up time (e.g., average and total amount), if appropriate to the study design | n/a |
| **Outcome data** | 15* | (a) Report outcomes as appropriate for the study design and summarize at all relevant levels of organization | 13-22 |
|  |  | (b) For proportions and rates, report the numerator and denominator |  |
|  |  | (c) For continuous outcomes, report the number of observations and a measure of variability |  |
| **Main results** | 16 | (a) Give unadjusted estimates and, if applicable, adjusted estimates and their precision (e.g., 95% confidence interval). Make clear which confounders and interactions were adjusted. Report all relevant parameters that were part of the model | 13-19 |
|  |  | (b) Report category boundaries when continuous variables were categorized | n/a |
|  |  | (c) If relevant, consider translating estimates of relative risk into absolute risk for a meaningful time period | n/a |
| **Other analyses** | 17 | Report other analyses done, such as sensitivity/robustness analysis and analysis of subgroups |  |
| **Key results** | 18 | Summarize key results with reference to study objectives | 1,22,30-31 |
| **Strengths and Limitations** | 19 | Discuss strengths and limitations of the study, taking into account sources of potential bias or imprecision. Discuss both direction and magnitude of any potential bias | 30-31 |
| **Interpretation** | 20 | Give a cautious overall interpretation of results considering objectives, limitations, multiplicity of analyses, results from similar studies, and other relevant evidence | 30-31 |
| **Generalizability** | 21 | Discuss the generalizability (external validity) of the study results | 31 |
| **Funding Transparency** | 22 | (a) Funding- Give the source of funding and the role of the funders for the present study and, if applicable, for the original study on which the present article is based | 32 |
|  |  | (b) Conflicts of interest-Describe any conflicts of interest, or lack thereof, for each author<br>(c) Describe the authors' roles- Provision of an authors' declaration of transparency is recommended<br>(d) Ethical approval- Include information on ethical approval for use of animal and human subjects<br>(e) Quality standards-Describe any quality standards used in the conduct of the research |  |

<sup>a</sup> Level of organization recognizes that observational studies in veterinary research often deal with repeated measures (within an animal or herd) or animals that are maintained in groups (such as pens and herds); thus, the observations are not statistically independent. This non-independence has profound implications for the design, analysis, and results of these studies.
<sup>b</sup> The word "participant" is used in the STROBE statement. However, for the veterinary version, it is understood that "participant" should be addressed for both the animal owner/manager and for the animals themselves.
\*Give such information separately for cases and controls in case-control studies and, if applicable, for exposed and unexposed groups in cohort and cross-sectional studies.

